# A double hit affecting the *IKZF1-IKZF2* tandem in immune cells of schizophrenic patients regulate specific symptoms

**DOI:** 10.1101/2024.09.09.612162

**Authors:** Iván Ballasch, Laura López-Molina, Marcos Galán-Ganga, Anna Sancho-Balsells, Irene Rodríguez-Navarro, Sara Borràs-Pernas, M Angeles Rabadan, Wanqi Chen, Carlota Pastó-Pellicer, Francesca Flotta, Joaquín Fernández-Irigoyen, Enrique Santamaría, Ruth Aguilar, Carlota Dobaño, Natalia Egri, Carla Hernandez, Miqueu Alfonso, Manel Juan, Jordi Alberch, Daniel del Toro, Belén Arranz, Josep M. Canals, Albert Giralt

**Affiliations:** Departament de Biomedicina, Facultat de Medicina, Institut de Neurociències, Universitat de Barcelona, 08036 Barcelona, Spain; Institut d’Investigacions Biomèdiques August Pi i Sunyer (IDIBAPS), 08036 Barcelona, Spain; Centro de Investigación Biomédica en Red sobre Enfermedades Neurodegenerativas (CIBERNED), 28031 Madrid, Spain; ZeClinics SL. and ZeNeuroid SL. Barcelona, Spain; Proteomics Platform, Navarrabiomed, Hospital Universitario de Navarra (HUN), Universidad Pública de Navarra UPNA, IdiSNA, 31008 Pamplona, Spain; ISGlobal, Hospital Clínic – Universitat de Barcelona, Barcelona,Catalonia, Spain; CIBER Enfermedades Infecciosas (CIBERINFEC), Barcelona, Catalonia, Spain; Servei d’Immunologia, Hospital Clinic Barcelona (HCB) - CDB, Fundació Clínic de Recerca Biomèdica - IDIBAPS, Spain; Plataforma d’Immunoteràpia HSJD-HCB, Barcelona, Spain; Parc Sanitari Sant Joan de Déu, CIBERSAM, Barcelona, Spain; Production and Validation Centre of Advanced Therapies (Creatio), Faculty of Medicine and Health Science, University of Barcelona, 08036 Barcelona, Spain

**Keywords:** IL-4, CXCL10, CCL5, Neuronal networks, Ikaros, Helios, lymphocytes, mouse models, cognitive symptoms, negative symptoms

## Abstract

Schizophrenia is a complex multifactorial disorder and increasing evidence suggests the involvement of immune dysregulations in its pathogenesis. We observed that IKZF1 and IKZF2, classic immune-related transcription factors (TFs), were both downregulated in patients’ peripheral blood mononuclear cells (PBMCs) but not in their brain. We generated a new mutant mouse model with a reduction in Ikzf1 and Ikzf2 to study the impact of those changes. Such mice developed deficits in the three dimensions (positive-negative-cognitive) of schizophrenic-like phenotypes associated with alterations in structural synaptic plasticity. We then studied the secretomes of cultured PBMCs obtained from human patients and identified potentially secreted molecules, which depended on IKZF1 and IKZF2 levels, and that in turn have an impact on neural synchrony, structural synaptic plasticity and schizophrenic-like symptoms in *in vivo* and *in vitro* models. Our results point out that IKZF1-IKZF2-dependent immune signals negatively impact on essential neural circuits involved in schizophrenia.

## Introduction

Schizophrenia is composed of a myriad of heterogeneous symptoms that challenges a unified description of the disease. This represents a global burden, as schizophrenia ranks within the top 10 causes of disabilities in developed countries, with 24 million people affected worldwide (GHDx). Furthermore, due to its early age of onset of 16-25 years(McGrath et al., 2008; Saha et al., 2005), the disease results in more than 12 million disability-adjusted life years annually. Behavioral signatures of schizophrenia are broadly divided into three categories: (1) positive symptoms (delusions and hallucinations); (2) negative symptoms (lack of pleasure, social withdrawal), and (3) cognitive symptoms (impairment of short-term working memory, attention deficit). Current treatments only provide some symptomatic relief for positive symptoms whereas treatments for cognitive and negative symptoms remain elusive(Remington et al., 2016). Notably, cognitive and negative symptoms are responsible for a major proportion of the disability associated with schizophrenia and are more constant over time(Remington et al., 2016; Tamminga et al., 1998). Also, the current lack of insight into the disease is not only due to the heterogeneity of the symptomatology (Remington, 2018), but also due to the complexity of the causative factors underlying the disease, including genetic and immunological (Davis et al., 2016). Therefore, what is urgently needed are novel mechanistic insights into the molecular paths underlying the negative and cognitive symptomatology subtypes of the disorder.

Although schizophrenia has been classically observed as a brain disease, current evidence points towards a multi-system network underlying positive, negative, and cognitive symptoms (Chaudhry et al., 2020; van Mierlo et al., 2020). In this sense, the communication between the immune and central nervous systems has already been described to be altered in schizophrenia (Momtazmanesh et al., 2019). As illustrative example, it has been shown that lymphocyte invasion is higher in patients with prevalence of negative symptomatology(Busse et al., 2012) and that levels of inflammatory cytokines (TNF-a and IL-6 or C-reactive protein) correlate with negative and cognitive symptoms in schizophrenia(Goldsmith et al., 2018; Johnsen et al., 2016; Liemburg et al., 2018). However, what we know so far is reduced to the description of circulating molecules that probably serve as a molecular bridge between these two systems (Rømer et al., 2023). Major mechanistic insights are still lacking, especially regarding the regulation of these molecules, their aberrant release, and the specific targeted neural circuits they affect. In this line, a highly promising target as a molecular bridge between the neural and immune system is the Ikaros family of transcription factors (TFs), which include Ikaros zinc finger 1 (*IKZF1* or Ikaros) and Ikaros zinc finger 2 (*IKZF2* or Helios). These TFs have been largely described to be crucial for the normal development of immune cells (T-cells, B-cells, and monocytes)(Georgopoulos et al., 1997) and their function (e.g. production of *mb-1*, *CD3*, *IL2, IL4,* and *NFkB* among others(Molnár and Georgopoulos, 1994)). Furthermore, evidence starts to emerge, that they could play an important role in schizophrenia, as they specifically affect the maturation of neuronal types primarily affected in the disease (medium spiny neurons, hippocampal pyramidal neurons, and cortical pyramidal neurons) (Alsiö et al., 2013; Giralt et al., 2020; Martín-Ibáñez et al., 2017; Martín-Ibáñez et al., 2010). Nevertheless, the function of the IKZF family in the communication between the immune and neural systems during the development of schizophrenia is practically unexplored.

In the present work, we hypothesized that immune alterations in schizophrenia could be associated with altered levels (and function) of *IKZF1* and *IKZF2*, and that these potential changes could impact the communication between the immune and central nervous systems during the progression of the disorder. To address this hypothesis, we first confirmed the altered state of the two transcription factors in human samples (brain and circulating immune cells). Subsequently, to validate their specificity and biological relevance, we used genetically modified mice devoid of *Ikzf1*, *Ikzf2*, or both.

Finally, to evaluate the aberrant communication between the immune and central nervous systems, we transferred the secretome of human circulating immune cells (PBMCs) into the brains of living mice. In these mice, we evaluated whether schizophrenia-like phenotypes were induced in terms of behavior and histological markers.

## Materials and methods

### Human post-mortem brain samples

The brain samples from schizophrenia (SCH) patients used in this study were provided by the Sant Joan de Déu Brain Bank (Sant Boi de Llobregat, Barcelona, Spain). The donation and obtention of samples were regulated by the ethics committee of the institution. The sample processing followed the rules of the European Consortium of Nervous Tissues: BrainNet Europe II (BNEII). All the samples were protected in terms of individual donor identification following the BNEII laws. Clinical diagnosis of SCH in donor subjects was confirmed premortem with DMS-IV (Diagnostic and Statistical Manual of Mental Disorders – 4th edition) and ICD-10 (International Statistical Classification of Diseases and Related Health Problems) criteria by clinical examiners. Most donors were hospitalized for more than 40 years and were re-evaluated every 2 years to monitor and update their clinical progression. Case information can be found in Supplementary Table 1. Control samples (hippocampus, putamen, and dorsolateral prefrontal cortex) were obtained from Banc de Teixits Neurològics (Servei Científico-Tècnics, Universitat de Barcelona, Barcelona, Spain) and the sample processing also followed the BNEII rules. Case information can be also found in Supplementary Table 1. All the procedures for the obtention of post-mortem samples followed the ethical guidelines of the Declaration of Helsinki and local ethical committees (Universitat de Barcelona: IRB00003099; Fundació CEIC Sant Joan de Déu: BTN-PSSJD).

### Immunoblotting

The tissue was lysed by sonication in 250 µl of lysis buffer as described elsewhere (Giralt et al., 2017). After lysis, samples were centrifuged at 12,000 r.p.m. for 20 min. Supernatant proteins (15 mg) from total brain region extracts were loaded in SDS– PAGE and transferred to nitrocellulose membranes (GE Healthcare, LC, UK).

Membranes were blocked in TBS-T (150 mM NaCl, 20 mM Tris-HCl, pH 7.5, 0.5 ml Tween 20) with 50 g l−1 non-fat dry milk and 50 g l−1 BSA. Immunoblots were probed with the following antibodies: anti-Ikzf1 (1:1000, D6N9Y, Cell Signaling Technology, Danvers, MA, USA,#14859), anti-Ikzf2 (1:1000, GeneTex, California. USA, #GTX115630). All blots were incubated with the primary antibody overnight at 4 °C by shaking in PBS with 0.2 g l−1 sodium azide. After three washes in TBS-T, blots were incubated for 1 h at room temperature with the corresponding horseradish peroxidase-conjugated antibody (1:2000; Promega, Madison, WI, USA) and washed again with TBS-T. Immunoreactive bands were visualized using the Western Blotting Luminol Reagent (Santa Cruz Biotechnology, #sc-2048) and quantified by a computer-assisted densitometer (Gel-Pro Analyzer, version 4, Media Cybernetics). For loading control, a mouse monoclonal antibody for actin was used (1:20,000; MP Biochemicals, #0869100-CF).

### Recruitment of human patients

The sample of this study (N◦PI17/00246, PI Belen Arranz) was recruited in the Outpatient clinic located in Cornellà, Barcelona, Spain (Parc Sanitari Sant Joan de Deu). Adult controls and patients with DSM-5 (Diagnostic and Statistical Manual of Mental Disorders) schizophrenia-spectrum disorder at any stage of the disease were included. The inclusion criteria for patients were: (1) adults over 18 years of age; (2) ability to speak Spanish correctly; and (3) signed informed consent. Exclusion criteria were: (1) history of head trauma with loss of consciousness and (2) organic disease with mental repercussions; (3) presence of an acute inflammatory process: (3.1) fever (>38◦C) or infection in the two weeks prior to the baseline interview, or (3.2) have received vaccinations in the past 4 weeks. This study was conducted following the ethical principles of the Declaration of Helsinki and Good Clinical Practice and the Ethics and Research Board from Parc Sanitari Sant Joan de Deu. All participants provided written informed consent prior to their inclusion in the study. Case information can be found in Supplementary Table 2.

### Human peripheral blood mononuclear cells (PBMCs) and isolation of CD4+ and CD8+ cells

Human peripheral blood mononuclear cells (PBMCs) were isolated from peripheral blood. The cell fraction corresponding to red blood cells and granulocytes (neutrophils, basophils, and eosinophils) was removed from whole blood by density gradient centrifugation as described elsewhere (Kleiveland, 2015). For this procedure, 10 ml blood samples were diluted with 10 ml of phosphate-buffered saline (PBS) pH-7.2. Consequently, 10 ml of diluted blood were placed in 15 ml tubes filled with 4ml of a density gradient medium with ρ□=□1.077 g/ml (e.g. Ficoll-Paque PLUS) and centrifuged at 500×g for 35 min at 20 °C without brake. PBMCs were then transferred to 50 ml tubes, the tubes were filled with PBS and centrifuged at 500×g for 10 min at 20 °C. PBMCs were then lysated for mRNA (see next section) or plated. For PBMCs plating, after elimination of the supernatant, PBMCs were resuspended in X-vivo medium (#BEBP02-055Q Lonza bioscience, Maryland, USA) supplemented with pen/strep 1%/1%, L-glutamine 1% and Hepes 0,02M. Cells were seeded in 24-well plates (1ml per well at 4· 10□ cells/ml) and left in an incubator at 37°C, 5% CO2. After 24 h, PBMCs were treated with PMA (50 ng/ml, #P1585, Sigma-Aldrich Chemical Co., St. Louis, MO, USA) and Ionomycin (1 µM, #I0634 Sigma-Aldrich Chemical Co.) for 6 h, then we centrifuged the cultures to take separately the supernatant and the pellet, both were stored at −80°C. For the CD4+ and CD8+ isolation the Pan T Cell Isolation Kit was used to isolate T cells from human PBMC previously obtained through negative selection following manufacturer’s instructions (Miltenyi, Cat. #130-096-535). T cells were then subjected to further magnetic labeling and separation to isolate CD4+ and CD8+ cells. First, T cells were incubated with CD8 MicroBeads (Miltenyi, Cat. #130-045-201) for 15 min at 4°C and cell suspension was applied onto the magnetic columns. The flow-through containing unlabeled cells was collected and considered the CD4+ enriched fraction. Then, labeled CD8+ cells were removed from the column with buffer. Both fractions were centrifuged at 300g x 7 min and the final cellular pellets were lysed with the proper buffer for subsequent RNA extraction.

### mRNA extraction and RT-qPCR

PBMC were homogenized and total RNA was extracted using PureLink RNA Micro Scale kit (#12183-016, Invitrogen) according to manufacter’s recommendations. RNA purity and quantity were determined with Nanodrop 1000 spectrophotometer (Thermo Fisher). 500 ng of purified RNA was reverse transcribed using the High Capacity cDNA Reverse Transcription Kit (Applied Biosystems, Cat. #436814). The cDNA synthesis was performed at 25°C for 10 min, at 37°C for 120 min and a final step at 85°C for 5 min in a final volume of 20 μl as instructed by manufacturer. Then, cDNA was analyzed by quantitative RT-PCR using PrimeTime qPCR Assays (Integrated DNA Technologies, Coralville, Iowa. USA). Human assays: IKZF1 (Hs.PT.58.25575505), IKZF2 (Hs.PT.58.2960172), and 18S (Hs.PT.39a.22214856.g). Mouse assays: IKZF1 (Hs.PT.58.25575505), IKZF2 (Hs.PT.58.2960172) and 18S (Hs.PT.39a.22214856.g).

Quantitative PCR was performed in 12 μl of final volume on 96-well plates using the Premix Ex Taq (Takara Biotechnology, #RR037A). Reactions included Segment 1: 1 cycle of 30 seconds at 95°C and Segment 2: 40 cycles of 5 seconds at 95°C and 20 seconds at 60°C. All RT-PCR assays were run in duplicate. To provide negative control and exclude contamination by genomic DNA, the PrimeScript RT enzyme was omitted in the cDNA synthesis step and samples were subjected to the PCR reaction in the same manner with each probe. RT-PCR data were quantified using the comparative quantitation analysis program of 64 MxProTM quantitative PCR software version 3.0 (Stratagene) and 18S gene expression was used as housekeeping gene. To analyze the relative changes in gene expression, the 2(–ΔΔ C(T)) method was used.

### Treatments with antipsychotics in mice

Female and male adult (12-week-old) wild type mice (C57BL/6J strain) were purchased from Jackson Laboratory (Cat. #000664). All mice (male and female, 50% each) were chronically (7 days) treated (i.p.) with vehicle or paliperidone (0.5mg/Kg dissolved in PBS, 5% DMSO) or clozapine (1mg/Kg dissolved in PBS, 5% DMSO). Doses were used following previous literature(MacDowell et al., 2017; Szlachta et al., 2017). One day after the end of the treatments all mice were sacrificed and blood samples were obtained by using a cardiac puncture as previously described(Houthuys et al., 2010). Then, peripheral blood mononuclear cells (PBMCs) were isolated as we did for humans. Blood pools from two mice were used to obtain enough PBMC mRNA concentrations.

### Animals

Ikzf1 deficient mice (Wang et al., 1996) were provided by Professor Katia Georgopoulos whereas Ikzf2 deficient mice were provided by Professor Philippe Kastner (Cai et al., 2009). Both lines, with C57BL/6 background, were backcrossed to obtain wild type (Ik^+/+^:He^+/+^), heterozygous for *Ikzf1* (Ik^+/-^:He^+/+^), heterozygous for *Ikzf2* (Ik^+/+^:He^+/-^) and double heterozygous mice (Ik^+/-^:He^+/-^). Adult males and females were used. We also used the Egr1-CreERT2 mice (Brito et al., 2022). These mice carry a bacterial artificial chromosome (BAC) including the Egr1 gene in which the coding sequence was replaced by that of CreERT2 fusion protein. They were crossed with R26RCE mice (Gt(ROSA)26Sortm1.1(CAG-EGFP)Fsh/Mmjax, Strain 004077, The Jackson Laboratory), which harbor the R26R CAG-boosted EGFP (RCE) reporter allele with a loxP-flanked STOP cassette upstream of the enhanced green fluorescent protein (EGFP) gene to create the double heterozygous mutant Egr1-CreERT2 x R26RCE mice for the experiments related with characterization of neural populations upon treatment with human secretomes. Single intraperitoneal injections of 4-hydroxytamoxifen (4-HT; #H7904, Sigma-Aldrich Chemical Co.) 50 mg/kg were administered to induce conditional Cre-dependent recombination. The animals were housed with access to food and water ad libitum in a colony room kept at 19–22 °C and 40–60 % humidity, under an inverted 12/12 h light/dark cycle (from 8 A.M. to 8 P.M.). All animal procedures were approved by local committees [Universitat de Barcelona, CEEA (136/19); Generalitat de Catalunya (DAAM 10786) following the European Communities Council Directive (86/609/EU).

### Behavioral tests

Behavioral phenotyping was performed following widely and previously detailed features of appropriate mouse models of schizophrenia (Powell and Miyakawa, 2006) addressing the three dimensions of the symptomatology. First, free exploration and then, evaluation of D-amphetamine-induced hyperlocomotion (D-amphetamine sulfate, 3 mg/kg; TOCRIS 2813), were both evaluated in the open field as described elsewhere (Sancho-Balsells et al., 2020). Next, sociability was assessed by using the three-chamber social interaction test as previously described (Sancho-Balsells et al., 2020). Finally, object recognition long-term memory was evaluated by performing the novel object recognition test (NORT) as we previously reported (Ballasch et al., 2023). In all behavioral studies animals were tracked and recorded with Smart junior 3.0 software (Panlab).

### Golgi staining

Fresh brain hemispheres were submerged and processed following the Golgi-Cox method as described elsewhere (Giralt et al., 2017). Two hundred microns sections were cut in 70 % EtOH on a vibratome (Leica) and washed in water for 5 min. Next, they were reduced in 16 % ammonia solution for 1 h before washing in water for 2 min and fixation in Na2S2O3 for 7 min. After a 2-min final wash in water, sections were mounted on superfrost coverslips, dehydrated for 3 min in 50 %, then 70, 80 and 100% EtOH, incubated for 5 min in a 2:1 isopropanol:EtOH mixture, followed by 1 × 5 min in pure isopropanol and 2 × 5 min in xylol. Bright-field images of Golgi-impregnated stratum radiatum dendrites from hippocampal CA1 pyramidal neurons or from pyramidal neurons of the medial prefrontal cortex (Layer V) or from medium spiny neurons in the striatum, were captured with a Nikon DXM 1200F digital camera attached to a Nikon Eclipse E600 light microscope (x 100 oil objective). Only fully impregnated neurons with their soma entirely within the thickness of the section were used. Image z-stacks were taken every 0.2 mm, at 1,024 × 1,024-pixel resolution, yielding an image with pixel dimensions of 49.25 × 49.25 µm. Segments of proximal dendrites were selected for the analysis of spine density. Only spines arising from the lateral surfaces of the dendrites were included in the study. Given that spine density increases as a function of the distance from the soma, reaching a plateau 45 µm away from the soma, we selected dendritic segments of basal dendrites 45 µm away from the cell body. The total number of spines was obtained using the cell counter tool in the ImageJ software.

### Quantitative proteomics

Proteins from secretome media were precipitated with acetone, and pellets dissolved in 8M Urea, 50mM DTT. Protein quantitation was performed with the Bradford assay kit (Bio-Rad, Hercules, CA, USA). Protein extracts (20ug) were diluted in Laemmli sample buffer and loaded into a 1,5 mm thick polyacrylamide gel with a 4% stacking gel cast over a 12.5% resolving gel. The run was stopped as soon as the front entered 3 mm into the resolving gel so that the whole proteome became concentrated in the stacking/resolving gel interface. Bands were stained with Coomassie Brilliant Blue, excised from the gel and protein enzymatic cleavage was carried out with trypsin (Promega; 1:20, w/w) at 37 °C for 16 h as previously described (Shevchenko et al., 2006). Purification and concentration of peptides were performed using C18 Zip Tip Solid Phase Extraction (Millipore). For LC-MS/MS, dried peptide samples were reconstituted with 2% ACN-0.1% FA (Acetonitrile-Formic acid) and quantified by NanoDropTM spectrophometer (ThermoFisher) before LC-MS/MS analysis using an EASY-1000 nanoLC system coupled to an Exploris 480 mass spectrometer (Thermo Fisher Scientific). Peptides were resolved using a C18 Aurora column (75µm x 25cm, 1.6 µm particles; IonOpticks) at a flow rate of 300 nL/min using a 60-min gradient (50oC): 2% to 5% B in 1 min, 5% to 20% B in 48 min, 20% to 32% B in 12 min, and 32% to 95% B in 1 min (A= FA, 0.1%; B = 100% ACN:0.1% FA). The spray voltage was set at 1.6 kV, and the capillary temperature at 275 °C. Sample data was acquired in a data-independent mode (DIA) with a full MS scan (scan range: 400 to 900 m/z; resolution: 60000; maximum injection time: 50 ms; normalized AGC target: 100%) and 25 periodical MS/MS segments, applying 20 Th isolation windows (0.5 Th overlap; resolution: 15000; maximum injection time: 22 ms; normalized AGC target: 3000%). Peptides were fragmented using a normalized HCD collision energy of 30%. Data were acquired in profile and centroid mode for full MS scan and MS/MS, respectively. For data analysis of quantitative proteomics DIA data files were analyzed using Spectronaut (v 17.3, Biognosys) by directDIA analysis (dDIA). Sample raw data was first processed by the Pulsar Spectronaut search engine to generate a spectral library. Pulsar then searched MSMS spectra against Uniprot-Swissprot isoforms Homo Sapiens database. MS1/MS2 calibration and main search tolerance were set to dynamic. The maximum precursor ion charge was set to 4, and fragment selection to intensity-based. Carbamidomethyl (C) was selected as a fixed modification, and Oxidation (M), Acetyl (Protein N-term), Deamidation (N), and Gln-> pyro-Glu as variable modifications (3 maximum modifications per peptide). The enzyme was set to trypsin in a specific mode (two missed cleavages maximum). The target-decoy-based false discovery rate (FDR) filter for PSMs, peptide, and protein groups was set to 1%. Once the Pulsar search was performed, the Spectronaut DIA search engine algorithm initiated the DIA search using the default settings (Proteotypicity filter=Protein group Specific) and filtering the precursors and protein groups by a 1% Q-value. The Perseus software (version 1.6.15.0) (Tyanova et al., 2016) was used for statistical analysis and data visualization. Identifications from the reverse database, common contaminants and proteins only identified through a modification peptide were removed. Label-free intensities were then logarithmized (base 2) and the samples were then grouped according to the experimental design. A filter of 70 % valid values at least in one group was applied to the resulting matrix followed by a logarithmic transformation (Log2). Then, data were imputated based on the normal distribution and normalized using the “Width adjustment” method. Statistical analysis was performed using “Two-sample tests” for the T-test of two selected experimental groups, based on permutation-based FDR statistics (250 permutations; FDR 1⁄4 0.07; S0 1⁄4 0.1). The resulting matrixes were exported containing the p-value (-Log) and fold-change (Log2) columns that, after its transformation into linear scale, were filtered by p<0.05 and 30% of significance and fold-change respectively.

### Luminex assay

We measured a panel of 30 cytokines, chemokines and growth factors in 14 supernatant samples by Luminex using the Cytokine Human Magnetic 30-Plex Panel LHC6003M from Life Technologies™. Briefly, 25 μL of the sample were tested by applying a modification to the manufacturer’s protocol by using half the volume of all reagents including the standards. This modification has been previously tested and showed no difference in assay performance compared to the original protocol and has been used in prior studies (Aguilar et al., 2019; Harding et al., 2022). Each plate included 16 serial dilutions (2-fold) of a standard sample with known concentrations of each analyte and two blanks. Samples were acquired on a Luminex® 100/200 instrument and analyzed with xPONENT® software 3.1. Concentrations of analytes were obtained by interpolating the median fluorescent intensity (MFI) to a 5-parameter logistic regression curve fitted with the R package (drLumi) (Sanz et al., 2017). If the algorithm did not converge, a 4-parameter log-logistic model was fitted. Limits of quantification (LOQ) were estimated based on the 30% threshold value for the coefficient of variation (CV) of the standard curve. Molecules outside the LOQ in more than 30% of samples were excluded from the study. For all other markers, observations outside LOQ were imputed. Results were reported as pg/mL.

### Hippocampal primary cultures and immunocytochemistry

Hippocampal neurons were obtained from E17.5 C57Bl/6 mice. The hippocampus was dissected and mechanically dissociated with a fire-polished Pasteur pipette. Cells were seeded at a density of 50,000 onto 12 mm coverslips placed in 24-well plates. Plates were previously precoated with 0.1 mg/mL poly-D-lysine (Sigma-Aldrich Chemical Co., St. Louis, MO, USA) and neurons were cultured in Neurobasal medium (Gibco-BRL, Renfrewshire, UK) supplemented with 1% Glutamax and 2% B27 (Gibco-BRL). Cultures were maintained at 37 °C in a humidified atmosphere containing 5% CO_2_. Primary neurons were fixed with 4% paraformaldehyde solution in PBS for 10 min and blocked in PBS-0.1 M glycine for 10 min. Then, cells were permeabilized in PBS-0.1% saponin 10 min, blocked with PBS-Normal Horse Serum 15% for 30 min, and incubated overnight at 4 °C in the presence of the following primary antibodies: mouse MAP2 (1:500, M1406, Sigma-Aldrich Chemical Co.) and rabbit PSD-95 (1:500, Cell Signaling, Ref: 3450S). Fluorescent secondary antibodies: AlexaFluor 488 goat anti-rabbit (1:100) and/or Cy3 goat anti-mouse (1:100; both from Jackson ImmunoResearch, West Grove, PA, USA) were incubated for 1 h at RT. Nuclei were stained with DAPI-Fluoromount (SouthernBiotech, Birmingham, AL, USA). Immunofluorescence was analyzed using a Leica Confocal SP5-II confocal microscope (Leica Microsystems CMS GmbH, Mannheim, Germany). Images were taken using an HCX PL APO lambda blue 63.0 × 1.40 OIL objective with a standard pinhole (1 AU), at 1024 × 1024-pixel resolution, 0.2 µm thick, and 5.0 digital zoom.

### Tissue fixation and immunofluorescence

Mice were euthanized by cervical dislocation. Left hemispheres were removed and fixed for 72 h in 4 % paraformaldehyde (PFA) in PBS. 40 µm coronal sections were obtained using a Leica vibratome (Leica VT1000S). Next, free-floating sections were washed three times in PBS, treated with NH_4_Cl for 30 min, and washed again three times with PBS. Floating sections were permeabilized and blocked with PBS containing 3 % Triton X-100, 0.02 % Azide, 2 % BSA, and 3 % NGS (Ab buffer) for 1 h at RT. After three washes in PBS, brain slices were incubated overnight at 4 °C with anti-GFP (1:500, Synaptic Systems, 132006) and anti-Parvalbumin (1:1250, Swant, PV27). Sections were washed three times and incubated for 2 h at RT with fluorescent secondary antibody Alexa Fluor 488 or 647 (1:400; from Jackson Immuno Research, West Grove, PA, USA). Nuclei were stained with DAPI-Fluoromount (SouthernBiotech, Birmingham, AL, USA). Sections were analyzed using a two-photon confocal microscope (Leica SP5).

### Imaging analysis

To evaluate neuronal morphology by using the Sholl analysis we employed the protocol described elsewhere (de Pins et al., 2019). For the experiments evaluating PSD-95-positive puncta in primary cultures, we followed the protocol previously described (Giralt et al., 2017). For quantification of the number of activated neural ensembles and the number of parvalbumin-positive cells in the hippocampal CA1 from Egr1-CreERT2 x R26RCE mice, we used the procedure previously reported by our group (Sancho-Balsells et al., 2023).

### 3D Cell culture preparation

Tridimensional neurospheres networks were developed as previously described (Rabadan et al., 2022). Briefly, E18 C57BL/6JOlaHsd pregnant mice were euthanized by cervical dislocation. Hippocampus were dissected in Neurobasal (Gibco) ice-cold media and incubated in 0.25% Trypsin-EDTA (Gibco) at 37° C for 20 min, followed by 5□min DNAse I (1□μg/ ml; Sigma-Aldrich Chemical Co.) incubation at room temperature. Mechanical dissociation of the dissected hippocampus was performed by repeated pipetting with a fire-polished glass Pasteur pipette until a homogenous cell suspension was obtained. Cell viability was determined by Trypan Blue exclusion assay. The cell solution was then centrifuged at 150g for 10□min, and the supernatant was removed. The resulting cell pellet was resuspended in the culture media containing Neurobasal media, 2% B27, 1% N2, 0.5□mM Glutamate, and 1% Penicillin/Streptomycin (Gibco). Cells were infected with AAV7m8.Syn.GCaMP6s.WPRE.SV40 virus (Unitat de Producció de Vectors, Universitat Autonoma Barcelona) for neuronal GCaMP6s calcium sensor expression. The viral infection was made at 1:1000 dilution at the single-cell suspension stage, just before seeding. For neurospheres network cultures, 5.5×10^4^ cells were seeded in poly-dimethylsiloxane (PDMS) customized molds containing an 18 mm^2^ square well. PDMS molds were fabricated in UV resin 3D printed cast (Formlab 3+ laser 3D printer) and polymerized overnight at 90° C. Neurospheres cultures were incubated in culture media at 37° C and 5% CO2. Imaging experiments were performed using a wide-field fluorescence microscope (Inverted microscope Zeiss Axio Observer Z1) and ZEN software (Zeiss) was used for image acquisition. Time-lapse videos were recorded at 25 Hz for 5□min at 37□°C and 5% CO_2_ in culture media (Neurobasal media, 2% B27, 0.5□mM Glutamate, and 1% Penicillin/Streptomycin (Gibco)). Videos were acquired at DIV28. All computations and visualizations were done using the standard Python-based ecosystems for scientific computing and data visualization (NumPy, SciPy, Seaborn, SciKitLearn, Matplotlib). Statistical significance was plotted using the stat annotations package (https://github.com/trevismd/statannotations). Video data was extracted from .czi files. For neurosphere segmentation was done using the Otsu thresholding technique. A time series of mean intensity values of each neurosphere was extracted. The signal was denoised using a Butterworth filter. After normalization, peaks were detected (prominence (1-mean)/3, height 0.5) and the signal was binarized. Neurospheres without any detected activity were discarded from the subsequent analysis. From the binarized signal the following parameters were computed: mean activity rate, mean peak duration, and average pairwise Pearson correlation of neurospheres. Statistical significance between different conditions was determined at DIV28 by one-way ANOVA and Dunnett’s as *post-hoc* test. A subset of neurospheres were fixed and stained against Tuj1 (1:1000, #T8660; Sigma).

### Mini-osmotic pumps implantation

Mice were deeply anesthetized with isoflurane (2% induction, 1.5% maintenance) and 2% oxygen and placed in a stereotaxic apparatus for osmotic minipump (model 1004; Alzet, Palo Alto, CA, USA) implantation. A brain infusion kit (#0008663) was also used to deliver into the lateral (left) ventricle (0.1 mm posterior to bregma, ±0.8 mm lateral to the midline, and −2.5 mm ventral to the parenchyma surface) 0.11 μL per hour of supernatants obtained from cultured peripheral blood mononuclear cells/PBMC at a protein concentration of 13 ng/µl. Cannulas were fixed on the skull with Loctite 454 (from Alzet). Minipumps, previously equilibrated overnight at 37 °C in PBS, were implanted subcutaneously in the back of the animal. After recovery, the mice were isolated to prevent detachment of the cannulas.

### Statistics

All data are expressed as mean ± SEM. Statistical analysis was performed using the unpaired two-sided Student’s t test (95% confidence), one-way ANOVA with Tukey’s as *post hoc* tests, or two-way ANOVA with Bonferroni’s *post hoc* test as appropriate and indicated in the figure legends. Values of p < 0.05 were considered statistically significant. All experiments in this study were blinded and randomized. All mice bred for the experiments were used for preplanned experiments and randomized to experimental groups. Visibly sick animals were excluded before data collection and analysis. Data were collected, processed, and analyzed randomly. The experimental design and handling of mice were identical across experiments. Littermates were used as controls with multiple litters (3–5) examined per experiment.

## Results

### *IKZF1* and *IKZF2* expression levels are specifically downregulated in peripheral blood mononuclear cells in patients with schizophrenia

Since we have previously reported that mice full knockout for *Ikzf1* or *Ikzf2* could develop brain-related deficiencies (Alsiö et al., 2013; Giralt et al., 2020; Martín-Ibáñez et al., 2017; Martín-Ibáñez et al., 2010), we first aimed to assess IKZF1 and IKZF2 protein levels in different brain regions strongly implicated in the pathophysiology of schizophrenia, namely the hippocampus, dorsolateral prefrontal cortex, and putamen. We evaluated such protein levels in post-mortem brain samples from patients with schizophrenia and matched control patients (Suppl Table 1). We did not detect changes in IKZF1 and IKZF2 protein levels in either, the hippocampus (Fig. 1a-b) or the DLPFC (Fig. 1c-d) or putamen (Fig. 1e-f) when comparing samples form schizophrenic patients with samples from control subjects. The same results were observed when mRNA levels were evaluated (Suppl. Fig. 1). We next hypothesized that, since IKZF1 and IKZF2 in adult mammals are both more enriched in circulating immune (lymphocytes) cells than in neural tissues (John and Ward, 2011), see also in *The Human Protein Atlas*), it would be conceivable that potential changes could be easier to be detected there rather than in the brain. Thus, we isolated peripheral blood mononuclear cells (PBMCs), which are mostly ∼70-90% lymphocytes (Sen et al., 2018), from human controls and patients with schizophrenia (Suppl Table 2). A descriptive evaluation of these PBMCs showed no differences in terms of cell density for either, neutrophils or lymphocytes or monocytes when comparing control subjects with patients with schizophrenia (Suppl Table 3). We then extracted their mRNA and evaluated *IKZF1* and *IKZF2* mRNA levels by RT-qPCR. Interestingly, we detected that *IKZF1* (Fig. 1g) and *IKZF2* (Fig. 1h) were both significantly downregulated in PBMCs mRNA from patients with schizophrenia compared to matched controls. We then looked at isolated CD4+ and CD8+ cells from the same PBMCs. We found that *IKZF1* (Fig. 1i) and *IKZF2* (Fig. 1j) mRNA levels were both downregulated in CD4+ cells. In contrast, *IKZF1* (Fig. 1k) and *IKZF2* (Fig. 1l) mRNA levels were normal in CD8+ cells. Next, to rule out the possibility of unspecific effects mediated by the medications on these changes we decided to chronically treat wild type mice with the two most common administered antipsychotics in our cohort of patients; paliperidone and clozapine. We found that *Ikzf1* and *Ikzf2* mRNA levels in mouse PBMCs were not affected by such treatments (Suppl Fig. 2). Altogether, these results indicate that the double reduction of *IKZF1* and *IKZF2* in PBMCs is localized in CD4+ cells. These alterations could play a role in the dysfunctions described for this cell type in the context of schizophrenia(Corsi-Zuelli et al., 2021).

**Figure 1.**
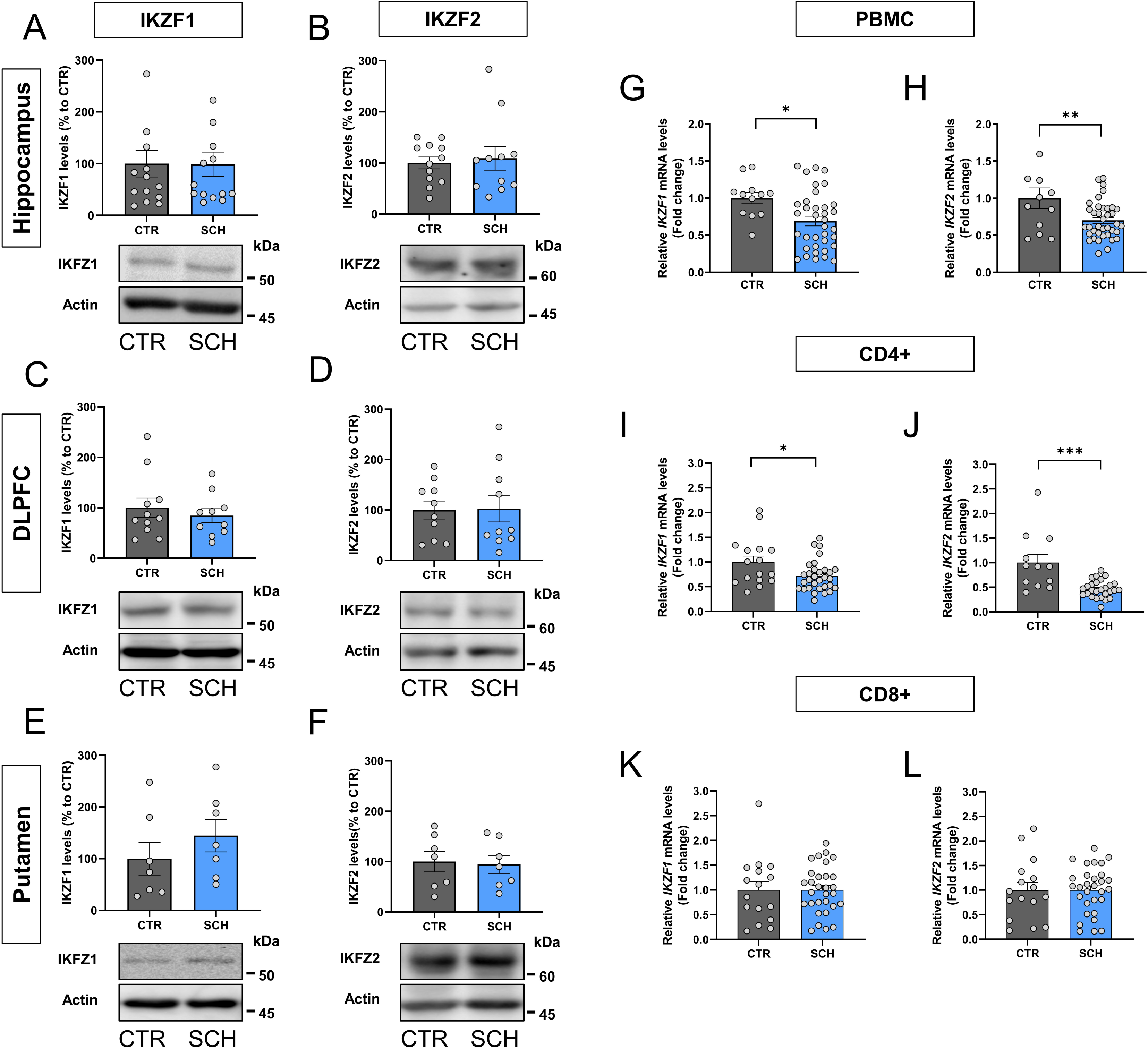
IKZF1 and IKZF2 levels in the brain and circulating immune cells of patients with schizophrenia. (**a**) Immunoblotting of total IKZF1 and (**b**) IKZF2, and actin as a loading control in the hippocampus of patients with schizophrenia (SCH) or matched controls (CTR). Upper panels depict quantification, lower panels show representative bands. (**c**) Immunoblotting of total IKZF1 and (**d**) IKZF2, and actin as a loading control in the dorsolateral prefrontal cortex (DLPFC) of patients with schizophrenia (SCH) or matched controls (CTR). Upper panels depict quantification, lower panels show representative bands. (**e**) Immunoblotting of total IKZF1 and (**f**) IKZF2, and actin as a loading control in the putamen of patients with schizophrenia (SCH) or matched controls (CTR). Upper panels depict quantification, lower panels show representative bands. (**g**) Results from RT-qPCR of total *IKZF1* and (**h**) *IKZF2* mRNA levels in peripheral blood mononuclear cells (PBMCs) isolated from patients with schizophrenia (SCH) or matched controls (CTR). (**i**) Results from RT-qPCR of total *IKZF1* and (**j**) *IKZF2* mRNA levels in CD4+ cells isolated from PBMCs in **i**-**j**. (**k**) Results from RT-qPCR of total *IKZF1* and (**l**) *IKZF2* mRNA levels in CD8+ cells isolated from PBMCs in **i**-**j**. Data are means ± SEM and they were analyzed using the two-tailed Student t-test. *p< 0.05, **p<0.01 and ***p<0.001 vs CTR.

### Mimicking the double *Ikzf1* and *Ikzf2* downregulation in mice results in a myriad of several schizophrenic-like disturbances

Although many different expressed genes (DEGs) and proteins (DEPs) have been identified in patients with schizophrenia, most of them have not been yet characterized or their role in the pathophysiology of the disease remain, at least, controversial (Rømer et al., 2023; Schizophrenia Working Group of the Psychiatric Genomics Consortium, 2014; Sullivan, 2013). The double reduction identified by us could not be an exception. Therefore, we aimed to characterize the relevance of this double reduction by mimicking it in mice. To do so we crossed heterozygous mice for *Ikzf1* (Ik^+/-^:He^+/+^, (Martín Ibáñez et al., 2010)) with heterozygous mice for *Ikzf2* (Ik^+/+^:He^+/-^, (Martín-Ibáñez et al., 2017)) to generate double mutant mice with a double reduction of both, *Ikzf1* and *Ikzf2* (Ik^+/-^:He^+/-^, Fig. 2a). Since both, adult (4-5 months of age) Ik^+/-^: He^+/+^ and Ik^+/+^:He^+/-^ mice displayed only punctual phenotypes compared to Ik^+/+^:He^+/+^ mice (Suppl. Fig. 3), henceforth we focused on the Ik^+/+^:He^+/+^ and Ik^+/-^:He^+/-^ genotypes (a.k.a. Wild type mice and double mutant mice) only. First, the appearance and body weight (Suppl. Fig. 4) were indistinguishable between the two groups, thus suggesting a normal health state in male and female Ik^+/-^:He^+/-^ mice. Then, to evaluate the potential schizophrenic-like phenotypes in our mutant mice, we performed a broad behavioral characterization aimed to evaluate positive, negative and cognitive symptomatology as previously established (Powell and Miyakawa, 2006). Regarding the positive-like phenotypes, we first observed that Ik^+/-^:He^+/-^ females (Fig. 2b) and Ik^+/-^:He^+/-^ males (Fig. 2c) displayed increased basal locomotion/agitation in the open field paradigm when compared with their matched Ik^+/+^:He^+/+^ controls. Furthermore, both, Ik^+/-^:He^+/-^ males (Fig. 2d) and females (Fig. 2e) showed an increased sensitivity to the D-Amphetamine in an open field arena. Regarding the negative-like phenotypes, we observed that in the three-chamber social interaction test there was a consistent alteration in sociability showed by both Ik^+/-^:He^+/-^ males (Fig. 2f) and females (Fig. 2g) compared with their matched controls. Finally, long-term declarative memory was also assessed by subjecting the groups of mice to the novel object recognition test (NORT). First, females Ik^+/-^:He^+/-^ were uncapable to recognize the new object with respect the old one when compared to Ik^+/+^:He^+/+^ controls (Fig. 2h). Similarly, Ik^+/+^:He^+/+^ males showed preference for the new object whereas Ik^+/-^:He^+/-^ males did not (Fig. 2i), suggesting the presence of detectable cognitive deficits in both Ik^+/-^:He^+/-^ males and females.

**Figure 2.**
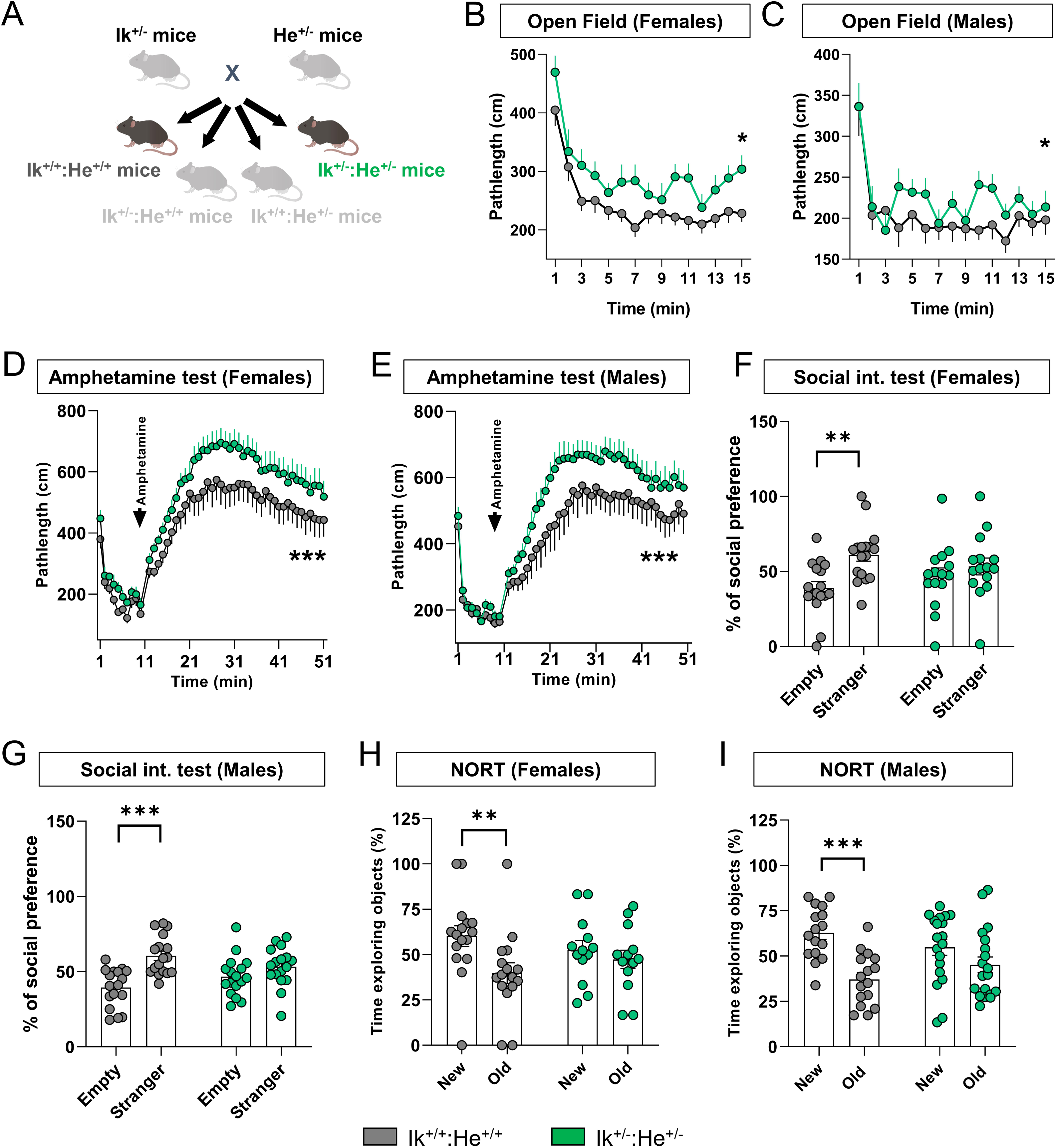
Generation and characterization of the Ik^+/-^:He^+/-^ double mutant mice. (**a**) Schematic representation of the adult mutant mice generated and used. Ik^+/-^ and He^+/-^ mice were crossed to generate the four genotypes; Ik^+/+^:He^+/+^, Ik^+/+^:He^+/-^, Ik^+/-^:He^+/+^ and Ik^+/-^:He^+/-^. Since the behavioral phenotype in Ik^+/+^:He^+/-^ and Ik^+/-^:He^+/+^ groups of mice were punctual compared to that in Ik^+/+^:He^+/+^ mice (Suppl. Fig. 3), we focused on the controls Ik^+/+^:He^+/+^ and the double mutant Ik^+/-^:He^+/-^ mice. Basal locomotor activity was evaluated in the two groups of mice separated into (**b**) females (Genotype effect: F_(1, 495)_ = 22,58, p < 0.0001) and (**c**) males (Genotype effect: F_(1, 538)_ = 9,518, p = 0.0021) in a 15 min testing session of free exploration in an open field. Induced agitation and sensitivity to the psychostimulant D-amphetamine was measured in a 45 min testing session upon injection of 5 mg/Kg of D-amphetamine in (**d**) female (Genotype effect: F_(49, 1450)_ = 20,37, p < 0.0001) and (**e**) male (Genotype effect: F_(48, 1666)_ = 21,28, p < 0.0001) mice from both groups. Sociability was evaluated in the three-chamber social interaction test (TCSIT). Mice from the two groups were subjected to the TCSIT and data were depicted for (**f**) females (Social preference effect: F_(1, 60)_ = 8,456, p = 0.0051) and (**g**) males (Social preference effect: F_(1, 64)_ = 19,36, p < 0.0001). Recognition memory was evaluated in the novel object recognition test (NORT). Mice from two groups were subjected to the NORT and data were depicted for (**h**) females (NORT preference effect: F_(1, 54)_ = 5,415, p = 0.0237) and (**i**) males (NORT preference effect: F_(1, 66)_ = 18,35, p < 0.0001). Data are means ± SEM and they were analyzed using the two-way ANOVA. *p< 0.05, ***p<0.001 vs Ik^+/+^:He^+/+^ mice in **b**, **d** and **e**. In **f**, **g**, **h,** and **i** data were analyzed using the two-way ANOVA with Bonferroni’s *post hoc* test; **p<0.01 and ***p<0.001 vs time exploring the stranger or the new object.

To further assess associated schizophrenic-like phenotypes such as dendritic spine loss in several brain regions as described elsewhere in post-mortem samples from human patients (Glausier and Lewis, 2013), we quantified spine density in hippocampal CA1 pyramidal neurons, in Medium Spiny Neurons (MSNs) of the dorsal striatum and in pyramidal neurons of the layer V in the medial pre-frontal cortex (mPFC) of Ik^+/+^:He^+/+^ and Ik^+/-^:He^+/-^ mice. First, we identified a significant reduction of dendritic spine density in the MSNs of both, Ik^+/-^:He^+/-^ males (Fig. 3a) and Ik^+/-^:He^+/-^ females (Fig. 3b) compared with their respective Ik^+/+^:He^+/+^ controls. In contrast, alterations in spine density of the CA1 pyramidal cells was only observed in Ik^+/-^:He^+/-^ males (Fig. 3c), but not in Ik^+/-^:He^+/-^ females (Fig. 3d). Finally, regarding to spine density in pyramidal neurons of the mPFC, no changes were observed in any condition (Fig. 3e-f). In conclusion, our newly generated double Ik^+/-^:He^+/-^ mutant mice mimic several features (behavioral and histological) in mice that are also observed in human patients with schizophrenia and models.

**Figure 3.**
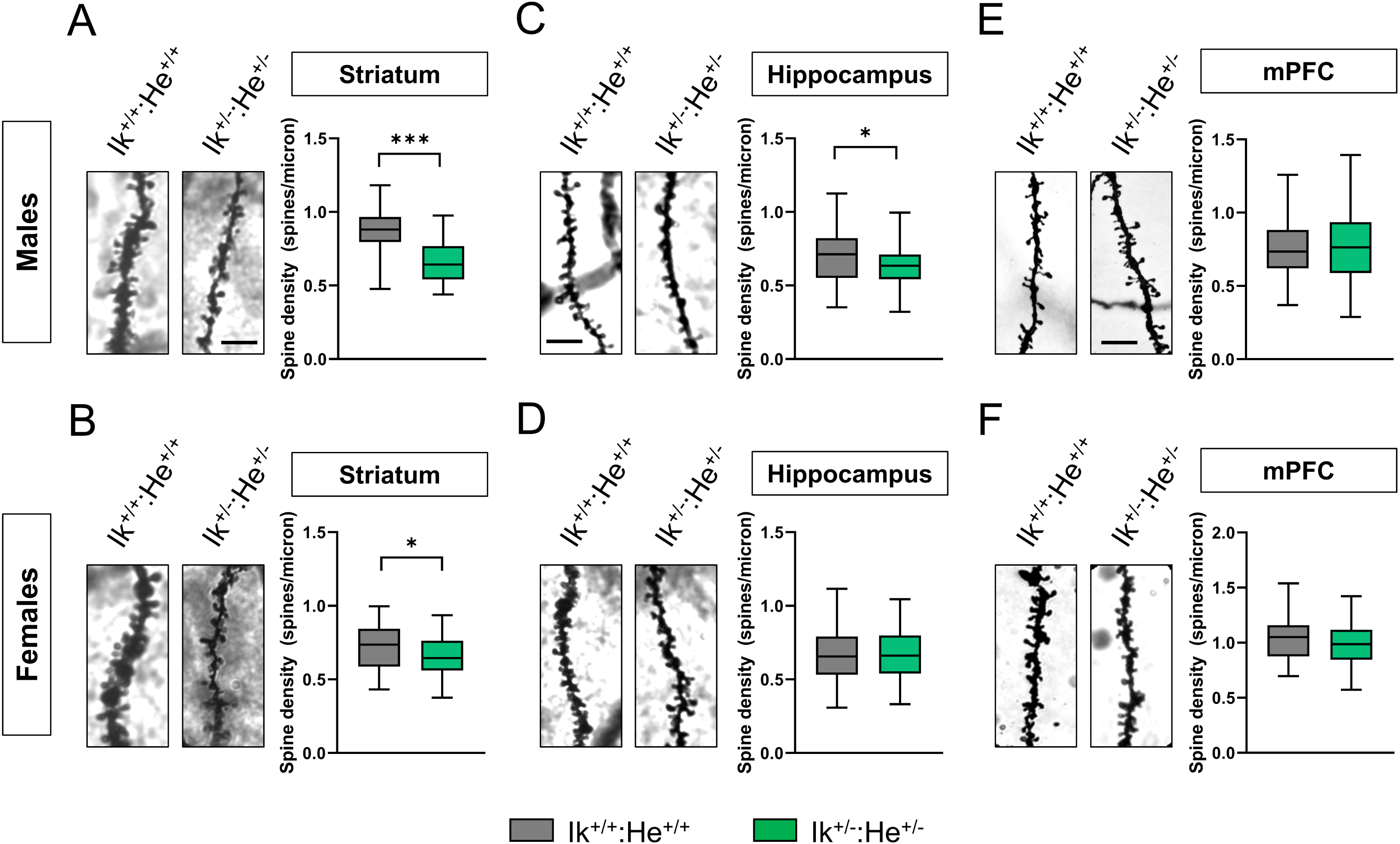
Characterization of structural synaptic plasticity in the Ik^+/-^:He^+/-^ mice. (**a**-**b**) Representative images (left panels) and quantification (right panels) of spine density in dendrites from medium spiny neurons (MSNs) of the dorsal striatum labeled with Golgi staining. Images were obtained in a bright-field microscope in adult (**a**) male and (**b**) female Ik^+/+^:He^+/+^ and Ik^+/-^:He^+/-^ mice. Scale bar, 5 μm. (**c**-**d**) Representative images (left panels) and quantification (right panels) of spine density in secondary apical dendrites from pyramidal neurons of the hippocampal CA1 labeled with Golgi staining. Images were obtained in adult (**c**) male and (**d**) female Ik^+/+^:He^+/+^ and Ik^+/-^:He^+/-^ mice. Scale bar, 5 μm. (**e**-**f**) Representative images (left panels) and quantification (right panels) of spine density in dendrites from pyramidal neurons of layer V in the frontal cortex labeled with Golgi staining. Images were obtained in adult (**e**) male and (**f**) female Ik^+/+^:He^+/+^ and Ik^+/-^:He^+/-^ mice. Scale bar, 5 μm. Data are means ± SEM and they were analyzed using the two-tailed Student t-test. *p< 0.05, ***p<0.01 vs Ik^+/+^:He^+/+^ mice. In **a**, n = 50–54 dendrites/genotype (from 7 mice/genotype). In **b**, n = 54–66 dendrites/genotype (from 7 mice/genotype). In **c**, n = 62–65 dendrites/genotype (from 7 mice/genotype). In **d**, n = 60–68 dendrites/genotype (from 7 mice/genotype). In **e**, n = 89–55 dendrites/genotype (from 7 mice/genotype). In **f**, n = 43–47 dendrites/genotype (from 7 mice/genotype). Scale bar in **a**, 5 µm.

### *IKZF1* and *IKZF2* levels in PBMC regulate the molecular profile of their secretome in patients with schizophrenia

Upon the observation that Ik^+/-^:He^+/-^ mice mimicked schizophrenia-like features, we next aimed to study which is the underlying molecular bridge between the circulating immune system and central nervous system that induces these phenotypes. We hypothesized that altered levels of *IKZF1* and *IKZF2* could impair the transcriptional activity of secreted molecules (secretome) by these circulating immune cells such as cytokines, chemokines, growth factors and other molecules. These impaired secretomes could, in turn, impair neural networks involved with the development of schizophrenia. To address these questions, we opted to use a more translational approach (Suppl. Fig. 5a). From patients with schizophrenia studied in figure 1g-h, we generated two sub-groups of patients; the first with normal levels of *IKZF1* and *IKZF2* (SCH^Ik+:He+^ group) in PBMC and the second with a double down-regulation of both, *IKZF1* (>40%) and *IKZF2* (>40%) in PBMC (SCH^Ik-:He-^ group) compared to control (CTR) patients (Suppl. Fig. 5b-c). All groups were matched in terms of number (n = 4-5/group), age (Suppl. Fig. 5d), sex, and the PANSS general score (Suppl. Table 4). Also, the number of circulating lymphocytes, neutrophils, platelets, and monocytes were indistinguishable between SCH^Ik+:He+^ and SCH^Ik-:He-^ groups (Suppl. Fig. 5e-h). We then obtained and cultured PBMC from all subjects and obtained their supernatants (a.k.a. secretomes; Suppl. Fig. 5a) containing the secretome of such cells. We first performed a protein expression screening in these secretomes using a mass spectrometry approach. First, principal component analysis (PCA) revealed that the two groups of patients with schizophrenia, SCH^Ik+:He+^ and SCH^Ik-:He-^, were different with respect to the CTR group (Fig. 4a). However, PCA indicated that SCH^Ik+:He+^ and SCH^Ik-:He-^ groups were slightly different between each other (Fig. 4a). Venn diagram representation indicated that 17 and 13 proteins were specifically altered in the SCH^Ik+:He+^ and SCH^Ik-:He-^ groups respectively compared to the CTR group. Also, only changes in 9 proteins were observed to be shared by both SCH^Ik+:He+^ and SCH^Ik-:He-^ groups when compared to CTR subjects (Fig. 4b-c). Proteins changing in this same sense included TUBA8, PFN1, TUBB, and DEFA1 (Fig. 4c-d). In contrast, specific altered proteins in the SCH^Ik+:He+^ group comprised ACTB, COTL1, FLNA, and PRTN3. On the other hand specific altered proteins in the SCH^Ik-:He-^ group involved ACTR3, SH3BGRL, LUZP1, CFL1, CCL5 (RANTES), C1RL and RHOC. To further complement these results from mass spectrometry we also performed a Luminex-based screen of several cytokines and chemokines in the supernatants from the three groups. Results indicated that some proteins were downregulated only in the SCH^Ik-:He-^ group compared to the CTR group such as GMSF and IL-4 (Fig. 4f) or MIP1A (Fig. 4g). Oppositely, CXCL10 was upregulated only in the SCH^Ik-:He-^ group compared to the CTR group (Fig. 4h). Finally, G-CSF and IL-10 were indistinguishable between groups (Fig. 4i). In summary, secretomes from SCH^Ik+:He+^ and SCH^Ik-:He-^ groups are significantly different between them and also compared to CTR secretomes.

**Figure 4.**
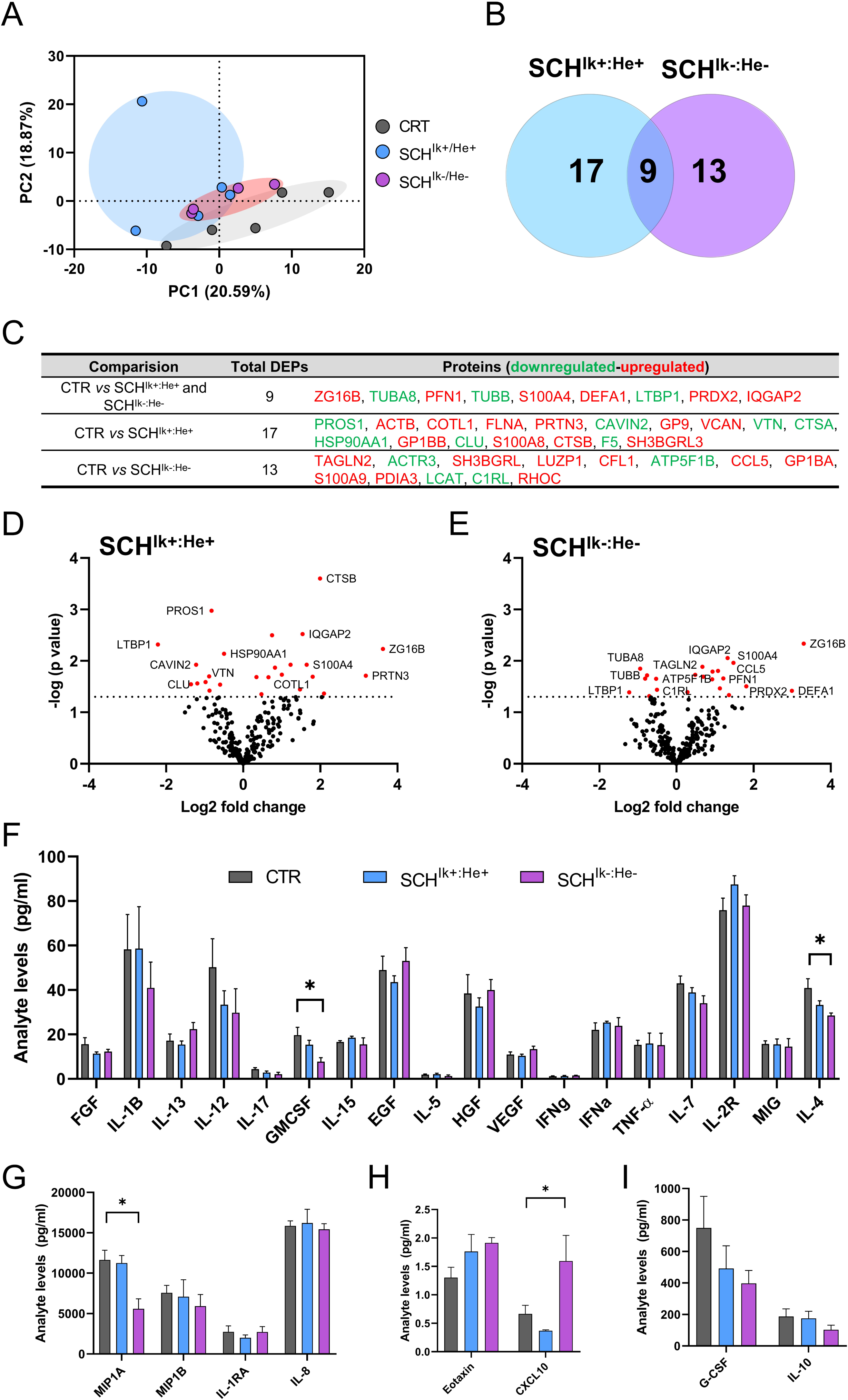
Molecular profile of PBMC secretomes from stratified patients with schizophrenia. From our results in figure 1g-h, we selected and stratified patients as follows: Patients with schizophrenia but with unaltered mRNA levels in PBMC of both, *IKZF1* and *IKZF2* respect to CTR subjects (SCH^Ik+:He+^ group; n = 5) and patients with a double reduction (≥40%) of *IKZF1* and *IKZF2* mRNA levels in PBMCs (SCH^Ik-:He-^ group; n = 4). These stratified patients were matched to control human subjects with normal *IKZF1* and *IKZF2* mRNA levels (CTR group; n=5). From these stratified patients, supernatants of cultured PBMC were collected and subjected to Mass Spectrometry. (a) Principal component analysis (PCA) for proteomics data from PBMC supernatants. The PCA plot represents the 14 subjects from the three subgroups (CTR in grey, SCH^Ik+:He+^ in blue, and SCH^Ik-:He-^ in violet) that indicates subtle proteomics profile differences between such supernatants. (**b**) Venn diagram showing the total number of proteins differentially expressed or DEPs in each of the SCH subgroups. The common DEPs (9) comparing both groups (SCH^Ik+:He+^ and SCH^Ik-:He-^) with respect to the CTR group, the specific DEPs (17, blue) comparing SCH^Ik+:He+^ with CTR, and the specific DEPs (13, violet) comparing SCH^Ik-:He-^ with CTR are depicted. (**c**) Table showing the specific down-regulated (green) and up-regulated (red) DEPs in each comparison extracted from **b**. (**d**) Volcano plot depicting DEPs when comparing supernatants from the CTR and SCH^Ik+:He+^ groups. Dashed horizontal line shows the p values cutoff, and the two vertical dashed lines indicate down/up regulated proteins. Red points indicate significantly DEPs. Dark points indicate non-significant DEPs. (**e**) Volcano plot depicting DEPs when comparing supernatants from the CTR and SCH^Ik-:He-^ groups as we did for **d**. Analyte array measured by a Luminex assay depicting the levels in pg/ml of (**f**) FGF, IL-1B, IL-13, IL-12, IL-17, GMCSF, IL-15, HGF, VEGF, IFNg, IFNa, TNFa, IL-17, IL-2R, MIG, IL-4 and (**g**) MIP1A, MIP1B, IL-1RA, IL-8 and (**h**) Eotaxin, CXCL10 and (**i**) G-CSF and IL-10. The analytes are depicted in four (**f**-**i**) different graphs due to their huge variability in terms of the range of concentrations. Data are means ± SEM and they were analyzed using the one-way ANOVA and the Dunnet’s *post hoc* in **f**-**i**. *p< 0.05 vs CTR group.

### *IKZF1* and *IKZF2* levels in PBMCs from patients with schizophrenia modulate neuronal structural plasticity in primary mouse hippocampal neurons

We observed that the secretomes of both, SCH^Ik+:He+^ and SCH^Ik-:He-^ groups, displayed different molecular profiles between each other and also compared to CTR subjects. Next, we aimed to evaluate whether such changes could exert some impact on neuronal functions. To do so, we first set up the safety of using such secretomes from human subjects in mouse neuronal cultures. We first pooled the secretomes for each group. All three pools (CTR, SCH^Ik+:He+^, and SCH^Ik-:He-^) had a protein concentration of 0,13 µg/µl. We then added two different dilutions, 1:1 and 1:10 from the CTR secretome pool to mouse hippocampal primary neurons at day in vitro 7 (DIV7, Fig. 5a). We also used two additional controls in this experiment, namely a naïve hippocampal primary culture without any treatment and a culture only treated with PBMC media X-Vivo (as a vehicle). We observed that 24 h after treatments, the dilution 1:1 induced neuronal degeneration *per se* whereas the dilution 1:10 did not affect neuronal morphology as compared with the two other controls (Fig. 5b and d). Thus, we used the 1:10 concentration henceforth for the ulterior experiments using the three secretome pools due to its safety for neuronal viability. Next, we evaluated the effect of these secretomes in neuronal branching by treating neurons at DIV7. While CTR and SCH^Ik+:He+^ secretomes did not induce significant changes on neuronal morphology, the SCH^Ik-:He-^ secretome induced a substantial reduction in the neuronal dendritic complexity (Fig. 5c-d). To further deepen on these effects, we then repeated the experiment but adding the secretomes from the three groups (CTR, SCH^Ik+:He+^ and SCH^Ik-:He-^) to primary hippocampal neuronal cultures at DIV20 (Fig. 5e). Then, 24 h later we quantified the number of PSD-95-positive clusters (as an excitatory post-synaptic marker) in these cultures. We observed a significant reduction of PSD-95-positive clusters in primary cultures treated with the SCH^Ik-:He-^ secretome but not in primary cultures treated with the SCH^Ik+:He+^ secretomes compared with the CTR group (Fig. 5f-g). We concluded that the different molecular profile displayed by the SCH^Ik-:He-^ PBMC secretome was probably the responsible for these specific effects on structural synaptic plasticity.

**Figure 5.**
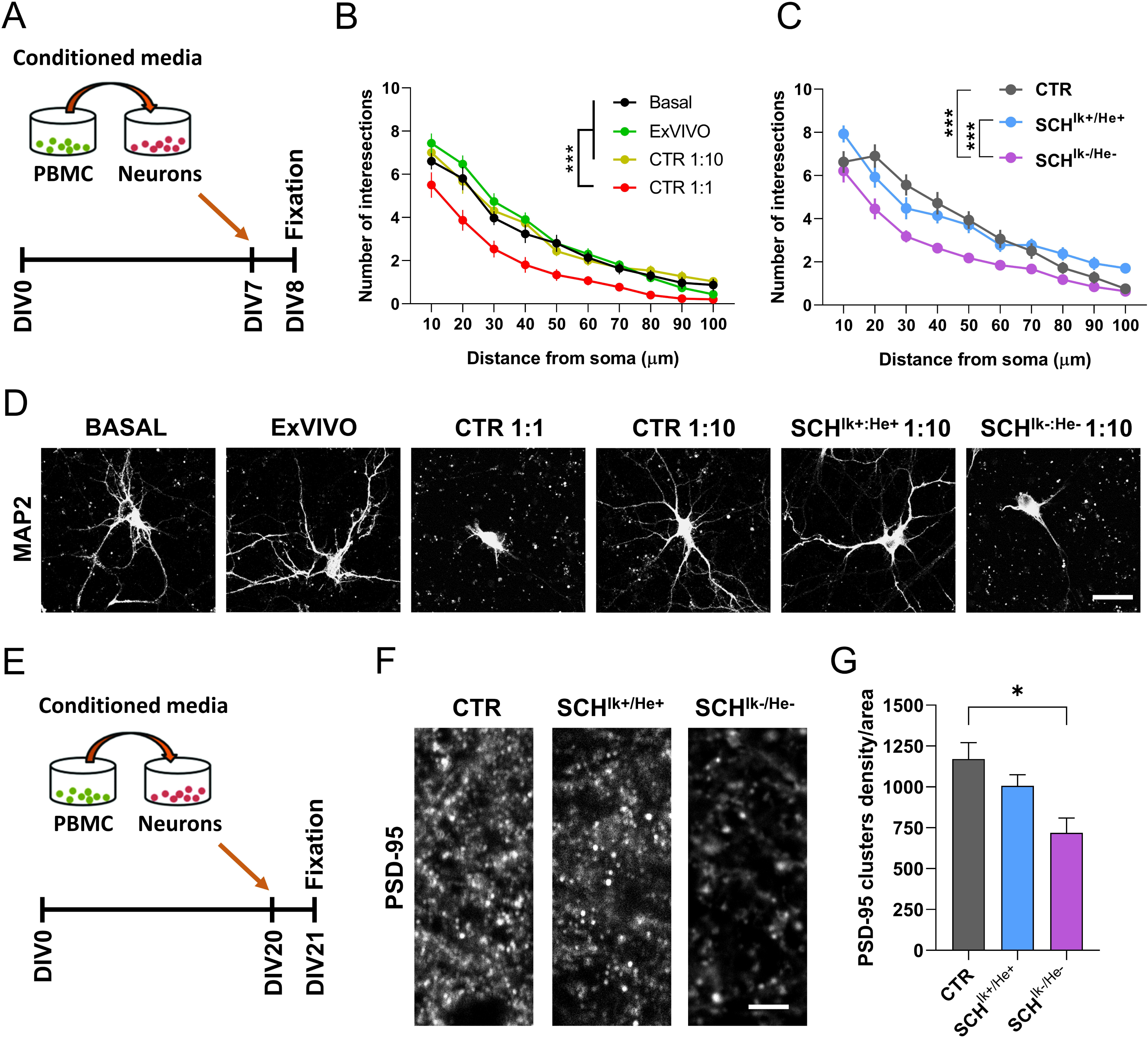
Effects of CTR, SCH^Ik+:He+^ and SCH^Ik-:He-^ supernatants in neuronal structural plasticity. (**a**) The experimental design is depicted. Supernatants (a.k.a. conditioned media) from cultured PBMC were added to primary hippocampal neurons at DIV7 and 24 h later, their dendritic morphology was assessed by using the Sholl analysis. (**b** and **d**) Different concentrations (1:1 and 1:10 supernatants from the CTR pool) and controls (X-Vivo media and naïve primary neurons) were employed (Group effect: F_(3, 1166)_ = 147,1, p < 0.0001). The 1:10 concentration was selected as the one to be used from **c** onwards because of its safety when compared with Basal and X-Vivo control conditions (**b** and **d**). (**c** and **d**) Effects of CTR, SCH^Ik+:He+^ and SCH^Ik-:He-^ supernatants on neuronal branching in primary hippocampal neurons using the selected (1:10) dose (Group effect: F_(2, 890)_ = 40,97, p < 0.0001). In **b**-**d** MAP2 staining was employed. (**e**) The experimental design to evaluate synaptic changes is depicted. Supernatants (conditioned media) from cultured PBMC were added to primary hippocampal neurons at DIV20 and 24 h later (**f**) the density of PSD-95 positive puncta per area was assessed in the three groups. (**g**) Quantification of PSD-95-positive puncta/area from **f** (F_(2, 9)_ = 6,905, p = 0.0152). Data are mean ± SEM. In **b** (n = 30 neurons/group) and **c** (n = 27-33 neurons/group) the two-way ANOVA was applied and Tukey’s multiple comparisons test was used as a *post hoc*. In **g** (n = 4 cultures/group) one-way ANOVA was applied, and Dunnett’s multiple comparisons test was used as a *post hoc*. Scale bar in **d**, 30 µm. Scale bar in **f**, 5 µm.

### The PBMCs secretome of patients with schizophrenia modulate neuronal dynamics: Role of *IKZF1* and *IKZF2*

Our previous results indicated that PBMC secretomes from patients with schizophrenia could play a role in structural synaptic plasticity, probably via modulation of synaptic components. To further explore this possibility, we benefit from a new approach called Modular Neuronal Network (MoNNets, (Rabadan et al., 2022)) designed to identify neural patterns (based on GcAMP6s-dependent calcium activity) in primary neurons resembling those observed in schizophrenia (Fig. 6a). Thus, MoNNets were generated, transduced with GcAMP6s and treated with CTR, SCH^Ik+:He+^ and SCH^Ik-:He-^ PBMC secretomes at DIV7, DIV14 and DIV21 (Fig. 6b). At DIV28 GcAMP6s-dependent calcium activity was recorded for 5 min in these MoNNets (Fig. 6b-c). Our results showed clear changes in activity in both groups of MoNNets treated with SCH^Ik+:He+^ and SCH^Ik-:He-^ secretomes compared to MoNNets treated with CTR secretome (Fig. 6d). These visual changes in activity in the raster plot were translated to a strong reduction in the average pairwise correlation, a measure of neuronal synchrony (Rabadan et al., 2022), in those MoNNets treated with SCH^Ik+:He+^, or SCH^Ik-:He-^ secretomes (Fig. 6e), which mimics what is observed in schizophrenia (Uhlhaas and Singer, 2010). Interestingly, we also observed a moderate decrease in activity rate in MoNNets treated with the SCH^Ik+:He+^ secretome, an effect even more robust when MoNNets were treated with the SCH^Ik-:He-^ secretome (Fig. 6f). Finally, only MoNNets treated with the SCH^Ik-:He-^ secretome suffered a reduction in the mean peak duration compared with the CTR condition (Fig. 6g). We concluded that, although both secretomes, the SCH^Ik+:He+^ and the SCH^Ik-:He-^, induced a reduction in MoNNets functional parameters (mainly synchrony) the SCH^Ik-:He-^ was the one that triggered the most severe and global alterations.

**Figure 6.**
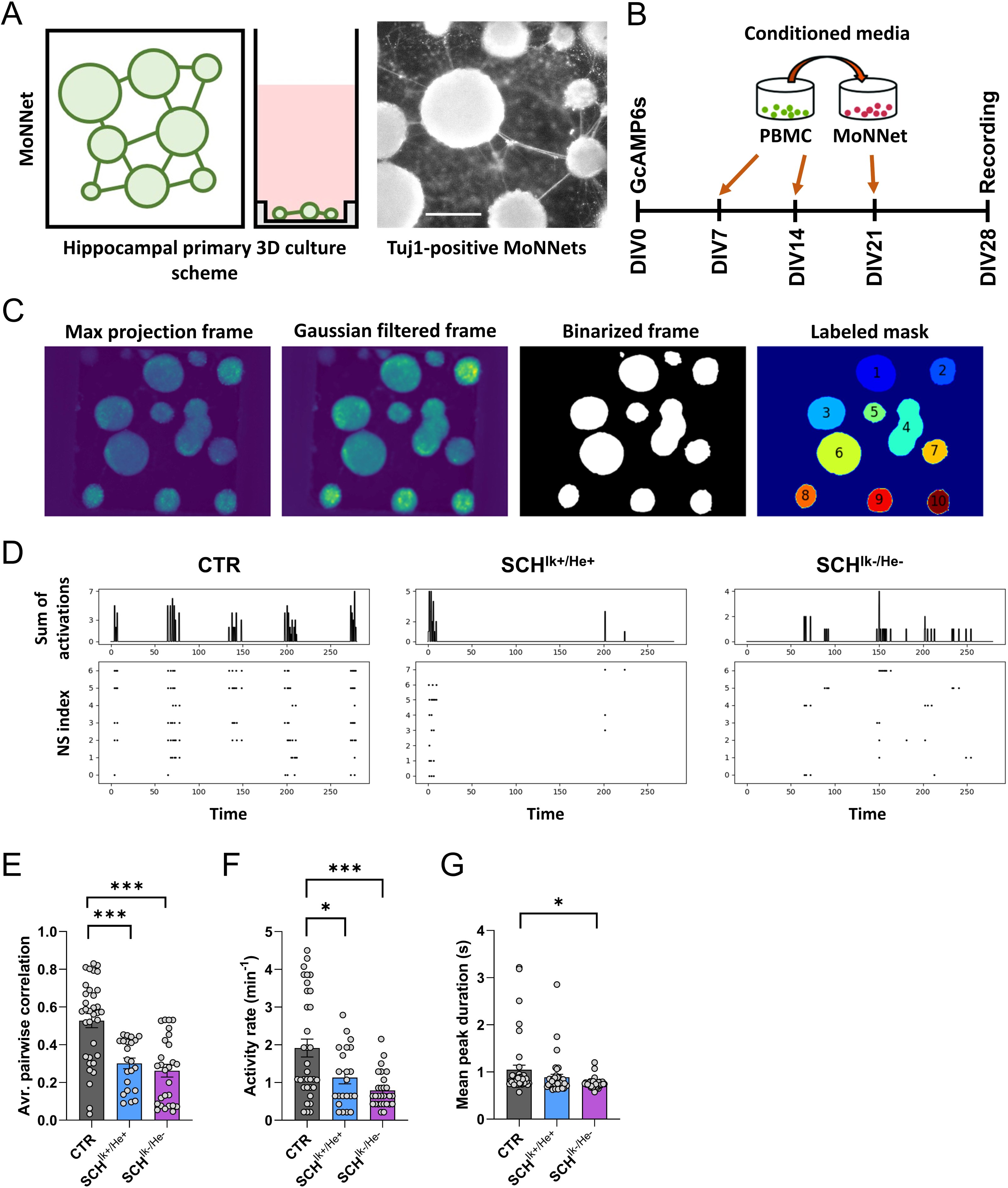
Impact of CTR, SCH^Ik+:He+^, and SCH^Ik-:He-^ supernatants in neuronal activity and synchrony. (**a**) Schematic overview of MoNNet approach (Left panel). Primary hippocampal cells were infected with AAV7m8.Syn.GCaMP6s.WPRE.SV40, and plated on a PDMS mold for self-organized assembly of MoNNet (a.k.a. neurospheres, Middle panel). Tuj-1 labeled MoNNets (right panel). Scale bar 250 µm. (a) The experimental design is depicted. Pooled supernatants (a.k.a. conditioned media) from each group (CTR, SCH^Ik+:He+^, and SCH^Ik-:He-^) of cultured PBMC were added to primary hippocampal neurons at DIV7, DIV14, and DIV21. At DIV28, their GcAMP6s-based activity was assessed. (**c**) Maxima projection showing GcAMP6s activity and its subsequent filtering (Gaussian > binarized > and final mask). System-wide cellular-resolution Ca^2+^ imaging was performed at 25 Hz. We analyzed the activity of each neurosphere and compared it with the rest of neurospheres. (**d**) Representative raster plots from DIV28 recordings in all three groups (CTR, SCH^Ik+:He+^, and SCH^Ik-:He-^). From the binarized signal the following parameters were computed: (**e**) average pairwise Pearson correlation of MoNNets (one-way ANOVA; F_(2, 80)_ = 18,31, p < 0.0001), (**f**) mean activity rate (one-way ANOVA: F_(2, 81)_ = 10,22, p < 0.0001) and (**g**) mean peak duration (one-way ANOVA: F_(2, 108)_ = 3,893, p = 0.023). In **e**-**g** one-way ANOVA was applied and the Dunnett’s multiple comparisons test was used as a *post hoc*. *p< 0.05 and ***p< 0.001 vs CTR.

### *IKZF1* and *IKZF2* levels in PBMCs from patients with schizophrenia modulate schizophrenic-like phenotypes with a specific impact on cognitive function

Our observations in primary neuronal cultures and in MoNNets suggested that neural networks could be influenced by an aberrant molecular profile in the PBMC secretome from patients with schizophrenia. Also, our results indicated that *IKZF1* and *IKZF2* levels modulate the severity of such impairments. To finally evaluate if such alterations have an impact in schizophrenic-like behavioral phenotypes and in neuronal networks *in vivo*, we used a mouse line that allows a permanent tagging of neurons activated by experience or relevant stimuli: the double mutant mice expressing Egr1-CreERT2 and R26RCE mice (Fig. 7a) (Brito et al., 2022; Longueville et al., 2021; Sancho-Balsells et al., 2023). We permanently and intraventricularly treated these transgenic mice with pooled secretomes from CTR, SCH^Ik+:He+^, and the SCH^Ik-:He-^ subjects using mini-osmotic pumps for 25 days (Fig. 7b-c) at a delivery ratio of 2,5 µl/day to the lateral ventricle. We performed a broad behavioral characterization of these mice assessing the three dimensions of schizophrenic-like phenotypes (Powell and Miyakawa, 2006) from day 5 to day 19 of the treatment. On day 20, mice were treated with 4-HT to induce recombination and permanent labeling of activated neural engrams due to the treatments with the secretomes. This design avoided the unspecific labeling of neural engrams due to the extensive behavioral characterization. Finally, brains were processed for histological studies on day 25 of treatment (Fig. 7c). Regarding the results, first, treated mice showed normal body weight (Fig. 7d) and appearance (data not shown) at the end of the study, suggesting that this treatment did not induce sickness-like states. Concerning the behavioral characterization, both SCH^Ik+:He+^ and the SCH^Ik-:He-^ groups showed no changes in agitation in the open field when compared with the CTR group (Fig. 7e). In contrast, both SCH^Ik+:He+^ and the SCH^Ik-:He-^ groups showed a clear and equal reduction in sociability compared to the CTR group in the three-chamber social interaction test (Fig. 7f). Finally, in the novel object recognition test, the SCH^Ik-:He-^ group but not the SCH^Ik+:He+^ group displayed specific alterations in novel object recognition memory in comparison with the CTR group (Fig. 7g). We then mapped the activation of potential aberrant engrams due to these secretomes in fixed brains from mice of the three groups. Since cognitive alterations seemed to be highly specific to the mice treated with the SCH^Ik-:He-^ secretome, we quantified the number of Egr1-dependent activated neural cells (GFP-positive) in the hippocampal CA1 (Fig. 7h) as suggested to be a core hippocampal sub-region in the physiopathology of schizophrenia (Wegrzyn et al., 2022). We observed that, although both groups, SCH^Ik+:He+^ and the SCH^Ik-:He-^ displayed an apparent aberrant activation of the Egr1-dependent activated ensembles, only the mice treated with the SCH^Ik-:He-^ secretome suffered a significant increase (Fig. 7h). Interestingly, increases on Egr1 levels have been previously reported in the auditory cortex (Iwakura et al., 2022) and in peripheral tissues of schizophrenic patients (Cattane et al., 2015). We then observed that altered structural synaptic plasticity was only detected in our mice treated with the SCH^Ik-:He-^ secretome. In particular, spine density in the CA1 pyramidal neurons (Fig. 7i) but not in medium spiny neurons of the dorsal striatum (Suppl. Fig. 6a) was increased in mice treated with the secretome from SCH^Ik-:He-^ patients compared to the CTR group. These differences, although a trend was observed, were not significant in mice treated with the SCH^Ik+:He+^ secretome. Such aberrantly increased number of Egr1-dependent activated ensembles correlated with a decrease in the number of parvalbumin-positive interneurons mostly observed in the SCH^Ik-:He-^ group, but also in the SCH^Ik+:He+^ group when compared with the CTR group (Fig. 7j). This reduction in parvalbumin interneurons could mediate the Egr1-dependent hyperactivation of CA1 pyramidal cells due to a lack of their inhibition (Valero and de la Prida, 2018). Overall, these results indicate that human secretomes from patients induce schizophrenic-like phenotypes in mice and that *IKZF1* and *IKZF2* levels in PBMC modulate the secretome conformation in a way that specifically impacts hippocampal activated neural ensembles likely through a reduction in parvalbumin interneurons function and ultimately impairing structural plasticity and associated cognitive skills.

**Figure 7.**
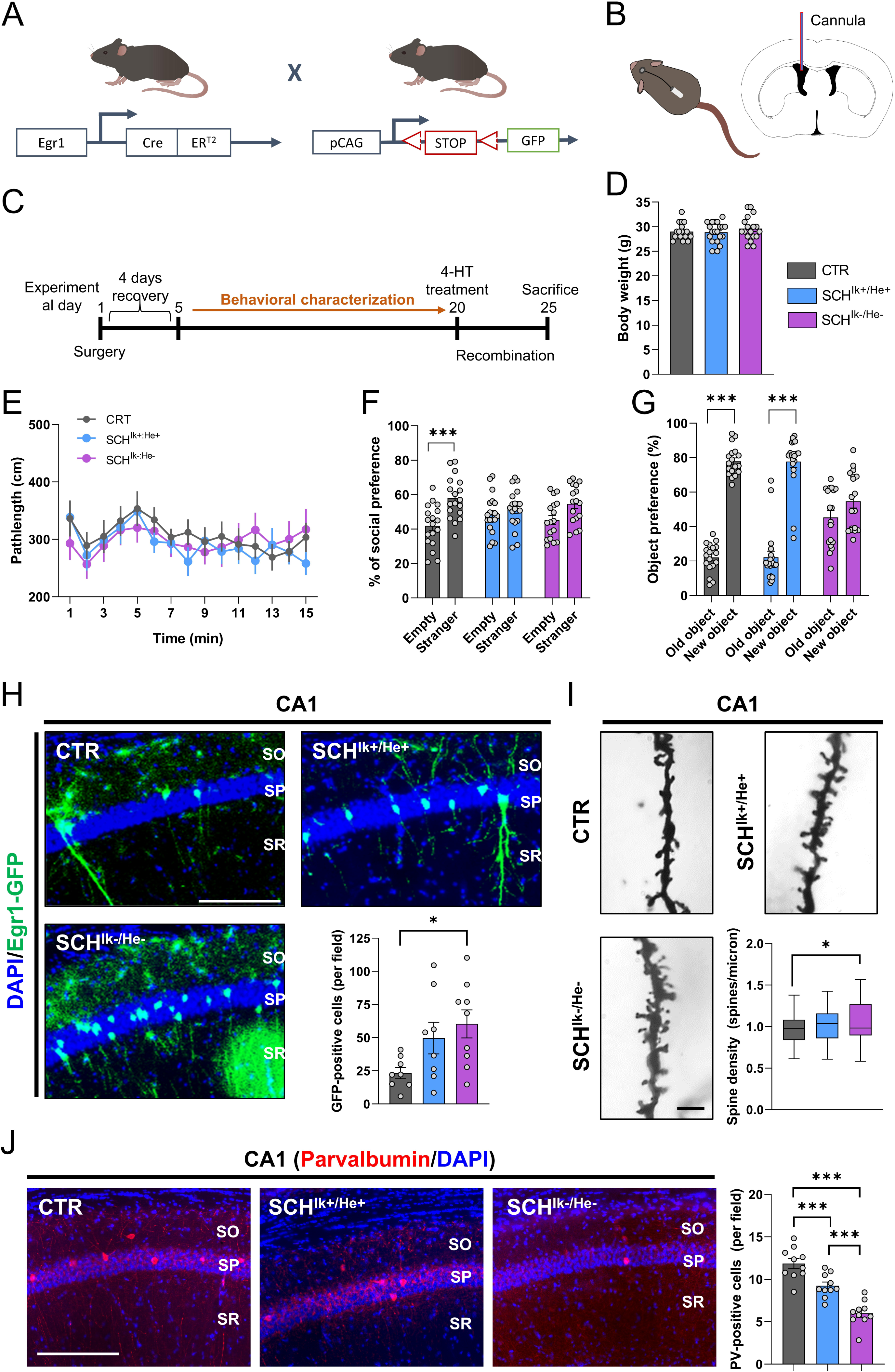
Schizophrenic-like phenotypes induced by the SCH^Ik+:He+^ and SCH^Ik-:He-^ supernatants when intraventricularly administered in mice. (**a**) Schematic representation of adult double-heterozygous-mutant Egr1-CreERT2 × R26RCE GFP mice used to label activated neural engrams upon treatment with supernatants. (**b**) Schematic representation of the intraventricular infusion of CTR, SCH^Ik+:He+^, and SCH^Ik-:He-^ supernatants. Mini-osmotic pumps infused 0,11 µl/h of supernatants at a concentration of 13 ng/µl of protein. (**c**) The experimental design is depicted. After surgical intervention, mice recovered for 4-5 days and then they were subjected to a broad behavioral characterization. At day 19, this behavioral characterization terminated and at day 20 mice were treated with 50mg/kg of 4-hydroxytamoxifen (4-HT) to induce recombination and labeling (GFP) of activated neural ensembles. At day 25, mice were processed to evaluate neural engrams formation and labeling and to evaluate spine density in the hippocampal CA1. (**d**) Body weight was monitored on the day of sacrifice. (**e**) Basal locomotion/agitation was evaluated by using the open field in the three groups of mice (CTR, SCH^Ik+:He+^, and SCH^Ik-:He-^). In the same groups of mice, (**f**) sociability was measured using the three-chamber social interaction test/TCSIT (Group effect: F_(1, 102)_ = 18,43, p < 0.0001). (**g**) The same groups of mice were also subjected to the Novel Object Recognition Test/NORT (Group effect: F_(1, 106)_ = 229,3, p < 0.0001). (**h**) Double fluorescent staining (DAPI in blue, GFP in green) in hippocampal CA1 from mice treated with one of each supernatant (CTR, SCH^Ik+:He+^, and SCH^Ik-:He-^). Graph (right-down) shows quantification of Egr1-dependent activated CA1 pyramidal cells (estimated number of GFP-positive cells/area of 500 µm^2^, Group effect: F_(2, 22)_ = 3,926, p = 0.034). Scale bar, 300 μm. (**i**) Representative images and quantification (right-down panel) of spine density in secondary apical dendrites of the CA1 pyramidal neurons labeled with Golgi staining (n = 72–121 dendrites/group from 7 mice/group). Images were obtained in a bright-field microscope in the three groups of mice treated with CTR, SCH^Ik+:He+^, and SCH^Ik-:He-^ supernatants (Scale bar, 5 μm). (**j**) Double fluorescent staining (left panels, DAPI in blue, parvalbumin in red) in hippocampal CA1 from the same mice as in **h**. Graph (right) shows quantification of parvalbumin-positive interneurons in the CA1 (estimated number of parvalbumin-positive cells/area of 500 µm^2^, Group effect: F_(2, 27)_ = 33,86, p < 0.001). Scale bar, 300 μm. Data are means ± SEM. In **d**, **h** and **I** one-way ANOVA was applied, and Tukey’s multiple comparisons test was used as a *post hoc*. In **e**, **f**, and **g**, two-way ANOVA was applied, and Bonferroni’s multiple comparisons test was used as a *post hoc.* *p< 0.05, vs CTR; ***p< 0.001, vs Empty or vs Old Object. CTR (n = 18); SCH^Ik+:He+^ (n = 20); SCH^Ik-:He-^ (n = 18). In **j** n = 10 in all groups.

## Discussion

In the present work, we show for the first time that both, *IKZF1* and *IKZF2*, are downregulated specifically in immune circulating cells, but not in the brains of patients with schizophrenia. We also observed in mice that a double *Ikzf1* and *Ikzf2* heterozygosis induced schizophrenic-like symptoms in the three dimensions (positive-negative-cognitive) of symptomatology. Exploring the secretome of these circulating immune cells, we observed that high-order cognitive functions are the most susceptible to be affected by the double *IKZF1* and *IKZF2* reduction. Specific and acutely hyperactivated hippocampal engrams, reduced number of parvalbumin interneurons and altered neuronal synchrony were associated with these phenotypes placing molecules such as IL-4 and CXCL10 as potential core signals mediating them.

Previous reports have already been addressed the molecular profile of PBMCs in patients with schizophrenia (Gardiner et al., 2013; Wu et al., 2016). Despite these previous studies and as far as we know, we show for the first time a double down-regulation of *IKZF1* and *IKZF2* levels. Interestingly, these and other studies have found that WNT signaling is a core pathway affected in these cells and that there is a correlation between WNT signaling and cognitive impairment in schizophrenia (Wu et al., 2016). In this line, the WNT family strongly regulates the IKZF genes in immune cells (Wu et al., 2022; Xiao et al., 2016), being a potential source of this double affectation of *IKZF1* and *IKZF2*. Regarding the cause of this double down-regulation, mutations in both *IKZF1* and *IKZF2* genes have never been associated with schizophrenia. As far as we know, no GWAS studies report associations between *IKZF1* and *IKZF2* point mutations and schizophrenia. In this context, we cannot rule out the possibility that, in some specific cases, rare mutations in these genes are affecting their levels in PBMCs and, in turn, impacting schizophrenia-related symptoms. However, a more plausible possibility is that alterations in *IKZF1* and *IKZF2* levels are indirectly provoked by other processes, such as immune and/or environmental challenges, as it has been proposed for the disorder(Ermakov et al., 2022).

To demonstrate the relevance of this double downregulation as a potential underlying molecular mechanism explaining the pathophysiology of schizophrenia, we have used both, a genetic and a translational model. In the genetic model, we have observed a highly schizophrenic-like phenotype (with phenotypes in all three dimensions, positive-negative-cognitive) in mice in terms of behavior and neuronal changes. However, although our data support the idea that an Ikzf1-Ikzf2 double reduction is necessary to display a full three-dimensional schizophrenic-like phenotype, it is worth mentioning that heterozygous Ik^+/-^:He^+/+^ and Ik^+/+^:He^+/-^ mice also display some specific schizophrenic-like phenotypes. This indicates that the observed phenotypes induced by the double reduction are partly synergistic but also partly additive. Regarding the translational model in which we used PBMC secretomes from patients, we concluded that an aberrant immune secretome with a double reduction of *IKZF1* and *IKZF2* from schizophrenic patients is enough to induce negative-like and cognitive-like symptoms. These results are in line with the idea that immune alterations are highly associated with such types of symptoms (Goldsmith and Rapaport, 2020; Malashenkova et al., 2021; Ribeiro-Santos et al., 2014). Noteworthy, the fact we did not observe differences in agitation in such translational mouse models could be because PBMC’ secretomes came from patients under treatment with antipsychotics. Although this statement is just speculative, it is in line with the large evidence stating that antipsychotics target principally positive symptoms whereas negative and cognitive symptoms remain largely unaffected (Malashenkova et al., 2021). Shall our hypothesis be true it would reinforce the fact of using agitation in mice as a positive-like phenotype/parameter.

Since our human PBMC secretomes induced so many schizophrenic-like phenotypes in terms of structural synaptic plasticity (Moyer et al., 2015), neuronal synchrony (Rabadan et al., 2022), and schizophrenic-like behaviors (Powell and Miyakawa, 2006), we then explored their molecular profile. We first found that levels of RANTES (CCL5), an upregulated cytokine in schizophrenia (Domenici et al., 2010; Frydecka et al., 2018), depended on *IKZF1* and *IKZF2* levels. However, the role of RANTES in schizophrenia is largely unknown. Another promising core molecule is IL-4. IL-4 rises as an interesting molecule coordinating the specific neuronal and cognitive symptoms observed in the induced SCH ^Ik-/He-^ *in vitro* and *in vivo* models. IL-4 is an attractive candidate since its levels are regulated by both, *IKZF1* and *IKZF2* in circulating immune cells (Gregory, 2006; Long et al., 2020; Quirion et al., 2009; Xie et al., 2021). Second, IL-4 regulates hippocampal-dependent memory (Brombacher et al., 2020; Gadani et al., 2012), and deficiency of its receptor, IL-4R, increased network activity (Hanuscheck et al., 2022) similar to what we observed in the hippocampus of our *in vivo* induced SCH ^Ik-/He-^ mouse model. In the context of schizophrenia, IL-4 has been shown to be significantly decreased in chronic schizophrenia-spectrum (Halstead et al., 2023). Furthermore, some single nucleotide polymorphisms (SNPs) in IL-4 have been related to the disorder (Schwarz et al., 2006). Additional studies have also shown that IL-4 levels are not related to positive psychotic symptoms (Fila-Danilow et al., 2012) but, instead, they correlate with negative symptoms (Şimşek et al., 2016). A third plausible candidate molecule underlying the defects observed in our SCH ^Ik-/He-^ mice is CXCL10 (IP10). Supporting this idea previous studies have observed enhanced neural activity accompanied by increased firing rate and excitability and alterations in synaptic network activity after chronic treatment with CXCL10 (Cho et al., 2009; Nelson and Gruol, 2004). These changes were sustained by reduced levels of GABA receptors, augmented levels of glutamate receptors and increased sensitivity of NMDA receptors (Cho et al., 2009; Nelson and Gruol, 2004). All these findings go in line with; the enhanced hippocampal Egr1-dependent activated ensembles in our SCH ^Ik-/He-^ mice, the basal hyperexcitability normally seen in schizophrenic patients and the reduced synchronized neural activity present in schizophrenic patients and in our SCH ^Ik-/He-^ *in vitro* MoNNet model. In this line, previous works (Grace and Gomes, 2019; Sonnenschein et al., 2020) have already proposed the hyperexcitability of the hippocampus as a core feature contributing to the three categories of symptoms seen in schizophrenia (Suppl Fig. 7a-c). We propose that our Ik^+/-^:He^+/-^ and SCH ^Ik-/He-^ models provide key underlying molecular clues, including IL-4 and CXCL10, supporting this model. As it has been reported in schizophrenic patients (Konradi et al., 2011), we also observed a reduction of parvalbumin-positive interneurons (PV+) in the hippocampal CA1 in our SCH ^Ik-/He-^ model. This reduction could be provoked by aberrant levels of IL-4 and CXCL10 from circulating PBMCs with decreased *IKZF1* and *IKZF2* levels. This altered secretome would induce a reduction in PV+ cells number and/or function since it has been shown that IL-4 reduces the risk of hyperexcitation (Hanuscheck et al., 2022) and CXCL10 over-expression reduces the levels of GAD65/67 (Cho et al., 2009) often accompanied of a ferroptosis processes (Liang et al., 2023). The reduced function of PV+ interneurons would provoke an hyperactivation of pyramidal neurons in the CA1 according to this circuit (Valero and de la Prida, 2018) and ultimately inducing the deficits in structural plasticity and impairments in cognitive function. However, although we provide evidences for this model, we cannot rule out other components allocated in these aberrant secretomes such as small molecules (RNAs) or extracellular vesicles (Alberro et al., 2021; Hunter et al., 2008). Future studies could address such possibilities.

Our study has some limitations, mostly concerning human samples. First, we used mRNA levels in human PBMCs to evaluate *IKZF1* and *IKZF2* expression; however, we do not have protein data. Additionally, there is a discrepancy in the number of recruited controls (CTR) and patients with schizophrenia (SCH) in the same experiments using PBMCs, which could introduce bias. Finally, we cannot rule out possible effects on neural *IKZF1* and *IKZF2* levels during earlier stages of the disorder since postmortem samples were collected from very old individuals, whereas PBMCs were collected from younger patients. In those same postmortem samples, there were other potential limitations such as differences in the postmortem time intervals. Therefore, although our results are supported by our studies using *in vivo* and *in vitro* models, the human data should be interpreted with caution.

Finally, our work fits with the idea that neural identities and probably function are both strongly modulated by peripheral signals and influences (López et al., 2007; Sun et al., 2022). One of them could be a double down-regulation of *IKZF1* and *IKZF2* levels in immune cells. In this sense, our results also have potential clinical implications. First, they could contribute to a better definition and stratification of patients based on their blood mRNA levels of *IKZF1* and *IKZF2*, which could be related to the severity of some symptoms. Additionally, it is tempting to speculate that different types of therapeutic strategies (drugs, RNA probes, Adeno-Associated Virus/AAVs) aimed at increasing mRNA levels of these transcription factors in circulating PBMCs could have positive therapeutic effects in patients with schizophrenia. Therefore, we strongly believe that future research should progress in the study of peripheral signals capable to play a role in the development of symptoms in schizophrenia and, probably, in other psychiatric affectations.

## Acknowledgements

We thank María Calvo from the Advanced Microscopy Service (Centres Científics i Tecnològics Universitat de Barcelona) for her help in the acquisition, analysis, and interpretation of the confocal images.

## Funding

This work was supported by grants from Ministerio de Ciencia e Innovación/ AEI/10.13039/501100011033/ and “FEDER” to A.G.: PID2021-122258OB-I00 and to J.A.: PID2020-119386RB-100. Our technician Ana López, was supported by a María de Maeztu Unit of Excellence (Institute of Neurosciences, University of Barcelona, CEX2021-001159-M, Ministry of Science and Innovation). The project that gave rise to these results received the support of a fellowship from “la Caixa” Foundation (ID 100010434). The fellowship code is LCF/BQ/DR21/11880016 to M.G.-G.

## Data availability

All data supporting the findings of this study are available within the paper and its Supplementary Information or it is available from the authors upon reasonable request.

## Ethics declarations

All the procedures for the obtention of post-mortem samples followed the ethical guidelines of the Declaration of Helsinki and local ethical committees (Universitat de Barcelona: IRB00003099; Fundació CEIC Sant Joan de Déu: BTN-PSSJD). The blood samples of this study (N◦PI17/00246, PI Belen Arranz) was recruited in the Outpatient clinic located in Cornellà, Barcelona, Spain (Parc Sanitari Sant Joan de Deu). All animal procedures were approved by local committees [Universitat de Barcelona, CEEA (136/19); Generalitat de Catalunya (DAAM 10786) following the European Communities Council Directive (86/609/EU).

## Declaration of competing interests

The authors declare no competing financial interests.

## Notes

### Competing Interest Statement

The authors have declared no competing interest.

