## Supplementary material for "A double hit affecting the *IKZF1-IKZF2* tandem in immune cells of schizophrenic patients regulate specific symptoms": Suppl figures and tables

**Supplementary figures and tables**

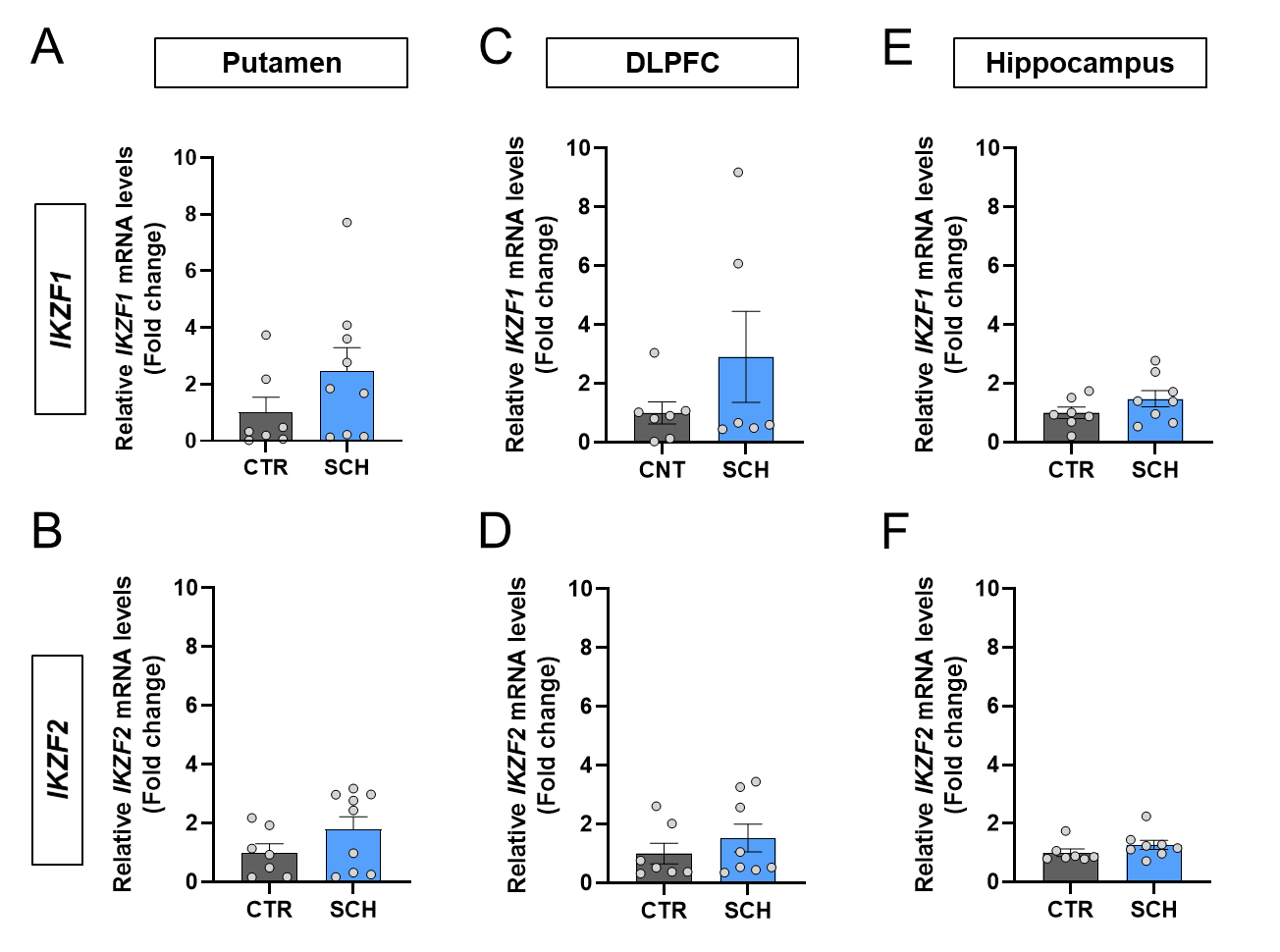

**Supplementary figure 1. Determination of** ***IKZF1* and *IKZF2* mRNA levels in different brain regions.** Results from RT-qPCR of total *IKZF1* and *IKZF2* mRNA levels in different brain regions namely putamen (**a** and **b** respectively), dorso-lateral prefrontal cortex (DLPFC, **c** and **d** respectively) and hippocampus (**e** and **f** respectively) from patients with schizophrenia (SCH) or matched controls (CTR). Demographics of the samples are displayed in supplementary table 1. Data are means ± SEM and they were analyzed using the two-tailed Student t-test. No differences were found in any comparison.

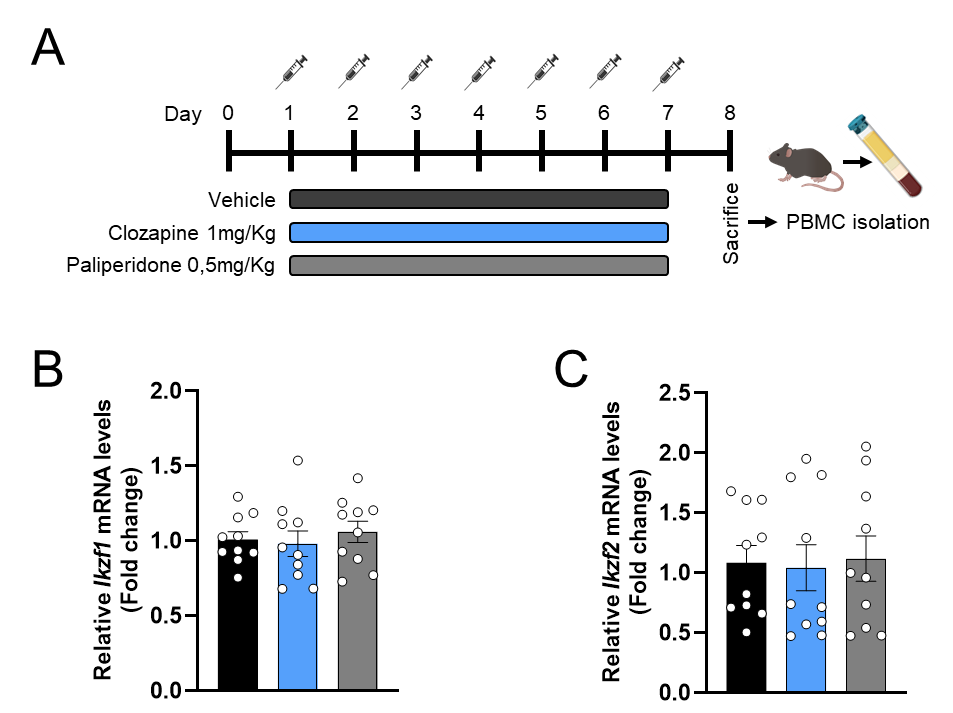

**Supplementary figure 2. Effects of antipsychotics on *Ikzf1* and *Ikzf2* mRNA levels in human PBMCs.** Timeline of experimental design (**a**). Mice were treated with vehicle (n = 20), 3 mg/Kg Clozapine (n = 20) or 0.5 mg/Kg Paliperidone (n = 20) i.p. for 7 days. On day 8, all mice were sacrificed and blood samples were obtained by cardiac puncture. Mice were pooled in groups of two to obtain enough peripheral blood mononuclear cells (PBMCs). PBMC were obtained and Ikzf1 (**b**) and Ikzf2 (**c**) mRNA levels were determined for each sample. The final “n” was 10/group. No differences were observed by treatments either in *Ikzf1* mRNA levels (one-way ANOVA, F_(2, 27)_ = 0,3250, p=0.725) or *Ikzf2* mRNA levels (one-way ANOVA, F_(2, 27)_ = 0,0471, p=0.954). Data are means ± SEM.

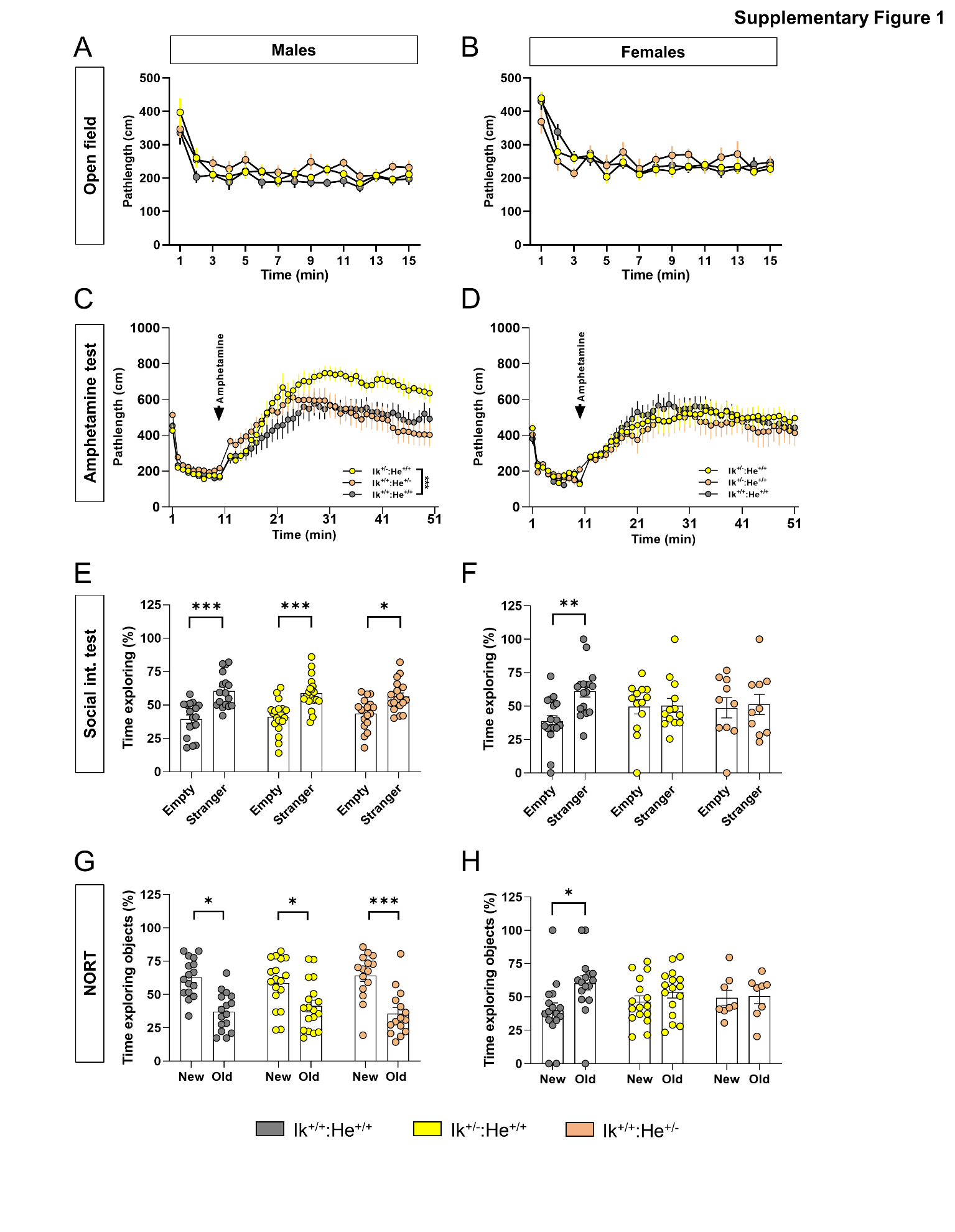

**Supplementary figure 3. Generation and characterization of the Ik^+/-^:He^+/+^ and Ik^+/+^:He^+/-^ mutant mice**. Basal locomotor activity was evaluated in the three groups of adult mice (Ik^+/+^:He^+/+^, Ik^+/-^:He^+/+^ and Ik^+/+^:He^+/-^ mice) separated in (**a**) males (Genotype effect: F_(2, 48)_ = 2.574, p = 0.087) and (**b**) females (Genotype effect: F_(2, 38)_ = 0.1940, p = 0.8244) in a 15 min testing session of free exploration in an open field. Induced agitation and sensitivity to the psychostimulant D-amphetamine was measured in a 45 min testing session in an open field upon injection of 5 mg/Kg of D-amphetamine in (**c**) adult male (Genotype effect: F_(2, 45)_ = 3.457, p = 0.0401) and in (**d**) adult female (Genotype effect: F_(2, 38)_ = 21,58, p = 0.8069) mice. Sociability was evaluated in the three-chamber social interaction test (TCSIT). Mice from the three groups were subjected to the TCSIT and data were depicted for (**e**) males (Social preference effect: F_(1, 102)_ = 53.58, p < 0.001) and (**f**) females (Social preference effect: F_(1, 74)_ = 3.482, p = 0.066). Recognition memory was evaluated in the novel object recognition test (NORT). Mice from the three groups were subjected to the NORT and data were depicted for (**g**) males (Novel object preference effect: F_(1, 92)_ = 48,68, p < 0.001) and (**h**) females (Novel object preference effect: F_(1, 74)_ = 4.296, p = 0.0417). Data are means ± SEM. Results were analyzed using the two-way ANOVA with Bonferroni’s *post hoc* test. ***p<0.001 vs Ik^+/+^:He^+/+^ mice in **c.** *p< 0.05, **p< 0.01 and ***p<0.001 vs time exploring the new object or vs time exploring the empty cage in **e**, **f,** and **g**. Male mice: Ik^+/+^:He^+/+^ (n = 16), Ik^+/-^:He^+/+^ (n = 19) and Ik^+/+^:He^+/-^ (n = 16). Female mice: Ik^+/+^:He^+/+^ (n = 16), Ik^+/-^:He^+/+^ (n = 8) and Ik^+/+^:He^+/-^ (n = 13).

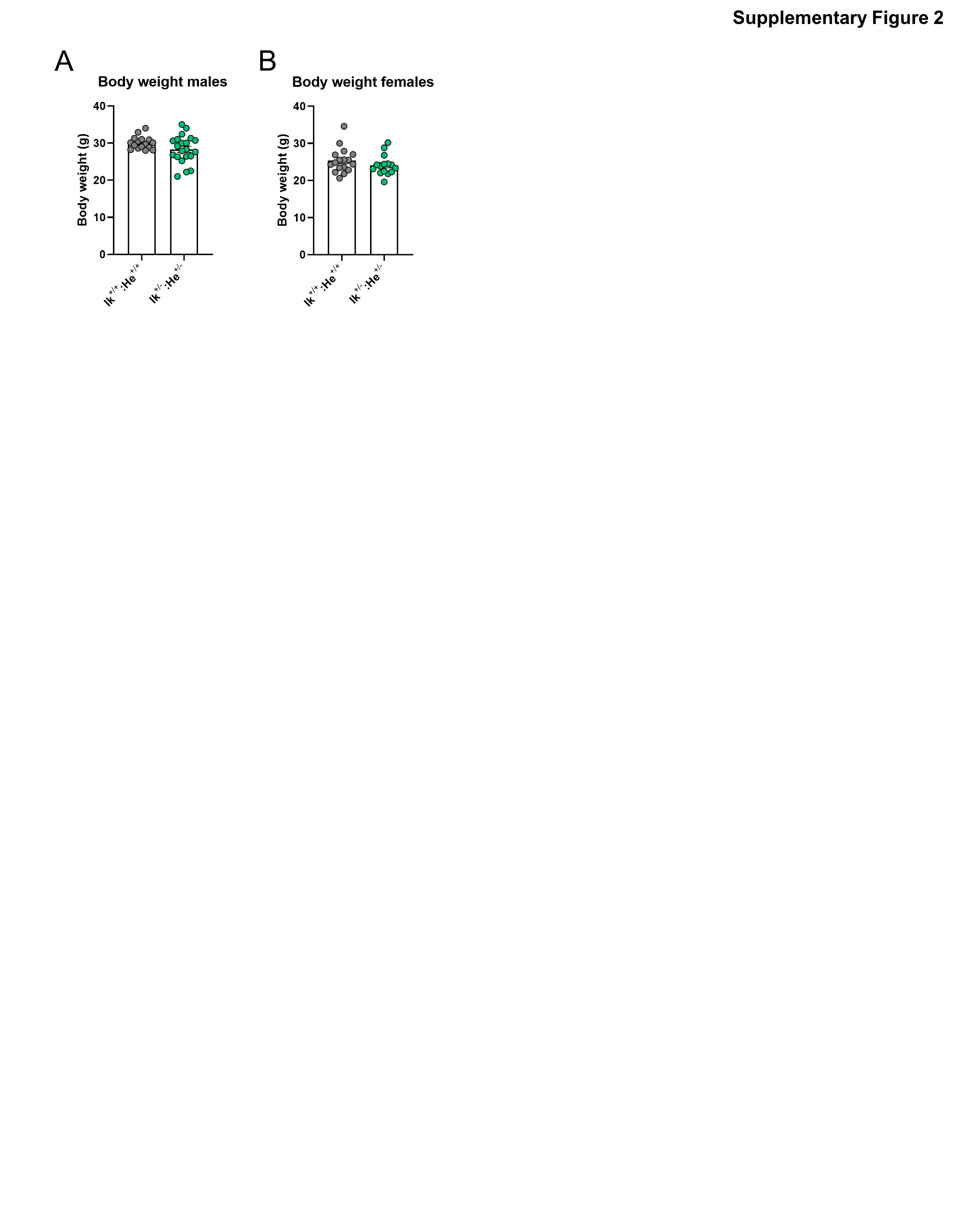

**Supplementary figure 4. Body weight of Ik^+/+^:He^+/+^ and Ik^+/-^:He^+/-^ mice**. (**a**) Body weight measured in adult males Ik^+/+^:He^+/+^ (n = 17) and Ik^+/-^:He^+/-^ (n = 21) mice. (**b**) Body weight measured in adult females Ik^+/+^:He^+/+^ (n = 17) and Ik^+/-^:He^+/-^ (n = 15) mice. Data are means ± SEM. Results were analyzed using the two-tailed Student t-test.

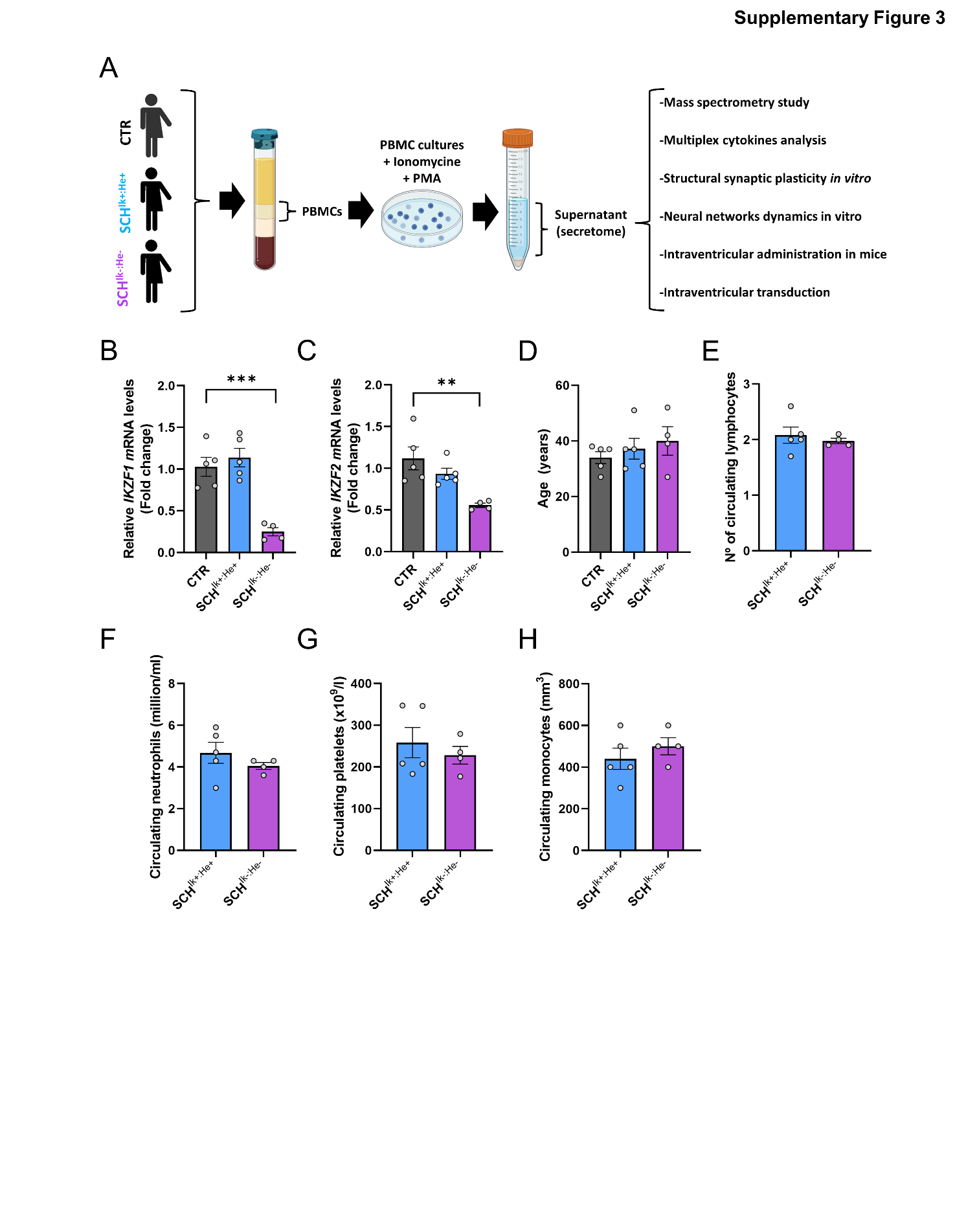

**Supplementary figure 5. Characterization of CTR, SCH^Ik+:He+^, and SCH^Ik-:He-^ supernatants.** (**a**) Schematic representation of the experimental design. First, patients were stratified according to whether they displayed significantly reduced (≥40% *IKZF1* and *IKZF2* mRNA levels (SCH^Ik-:He-^) or not (SCH^Ik+:He+^). These patients were matched with controls (CTR). Peripheral blood mononuclear cells (PBMCs) from the three groups of patients were isolated, cultured and stimulated (with Ionomycin and PMA) to obtain their secretomes (a.k.a. supernatants or conditioned media). Those supernatants were subjected to comprehensive characterization (Mass spectrometry, *in vitro* and *in vivo* studies, Multiplex cytokines analysis). As indicated, the three groups of patients were selected based on their psychiatric affectation (schizophrenia or not) and based on their (**b**) *IKZF1* and (**c**) *IKZF2* mRNA levels in PBMCs. Patients were matched by sex (**Suppl. Table 3**) and age (**d**). To determine that both groups, SCH^Ik+:He+^ and SCH^Ik-:He-^, are homogeneous and comparable, the number of circulating neutrophils (**f**), number of circulating platelets (**g**), and number of circulating monocytes (**h**) were proven to be similar. Data are means ± SEM. Results were analyzed using the one-way ANOVA with Dunnett’s *post hoc* test in **b**, **c** and **d**. Results in **e**, **f**, **g**, and **h** were analyzed using the two-tailed Student t-test. **p< 0.01 and ***p<0.001 vs CTR group**.** CTR (n = 5), SCH^Ik+:He+^ (n = 5) and SCH^Ik-:He-^ (n = 4).

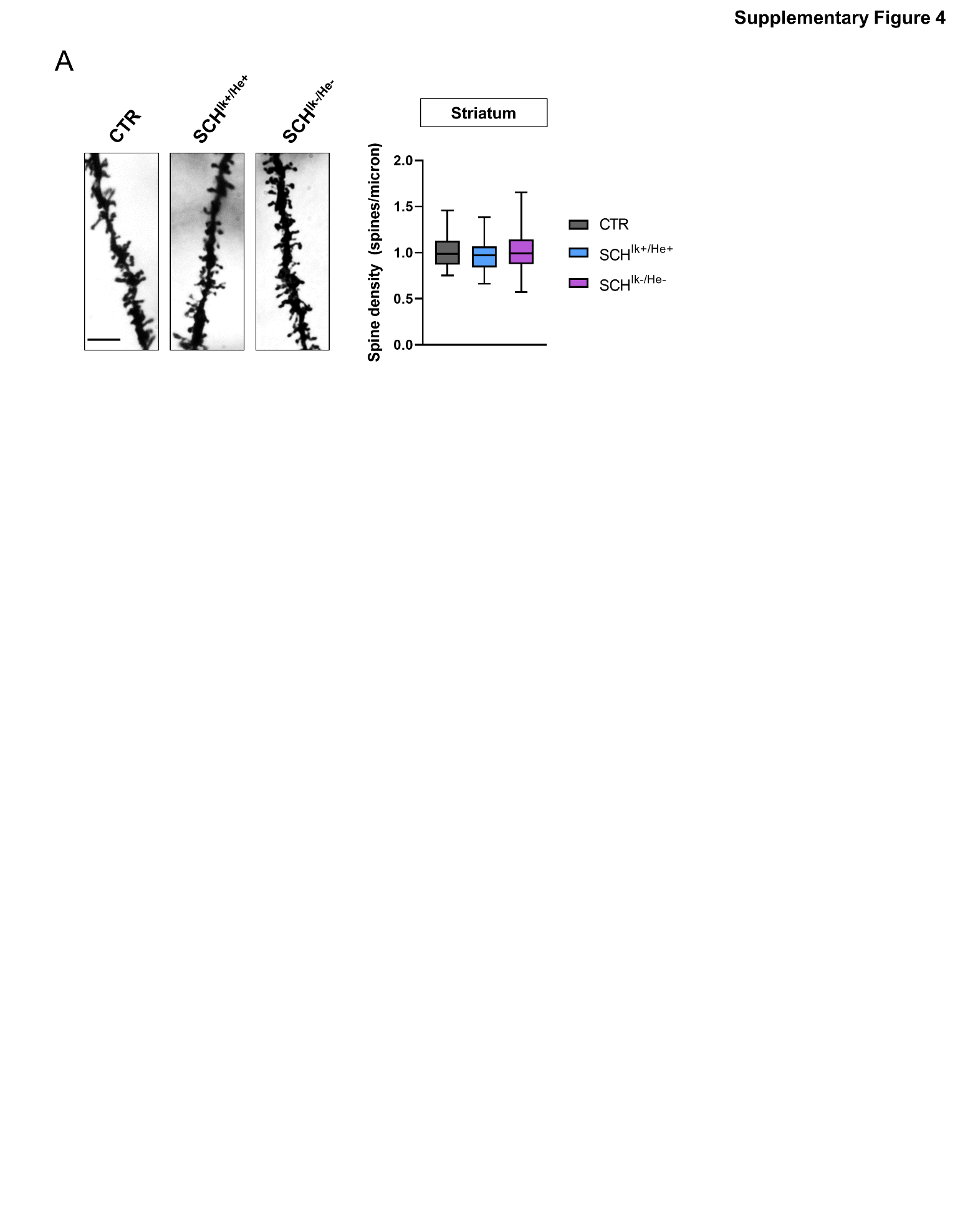

**Supplementary figure 6. Characterization of striatal structural synaptic plasticity in mice intraventricularly administered with CTR, SCH^Ik+:He+^ and SCH^Ik-:He-^ supernatants**. (**a**) Representative images (left panels) and quantification (right panels) of spine density in dendrites from medium spiny neurons (MSNs) of the dorsal striatum labeled with Golgi staining. Images were obtained in a bright-field microscope in adult male mice intraventricularly administered with CTR, SCH^Ik+:He+^ and SCH^Ik-:He-^ supernatants. Scale bar, 5 μm. Data are means ± SEM and they were analyzed using one-way ANOVA and Tukey's multiple comparisons test was used as a *post hoc*. N = 55–71 dendrites/genotype (from 7 mice/genotype).

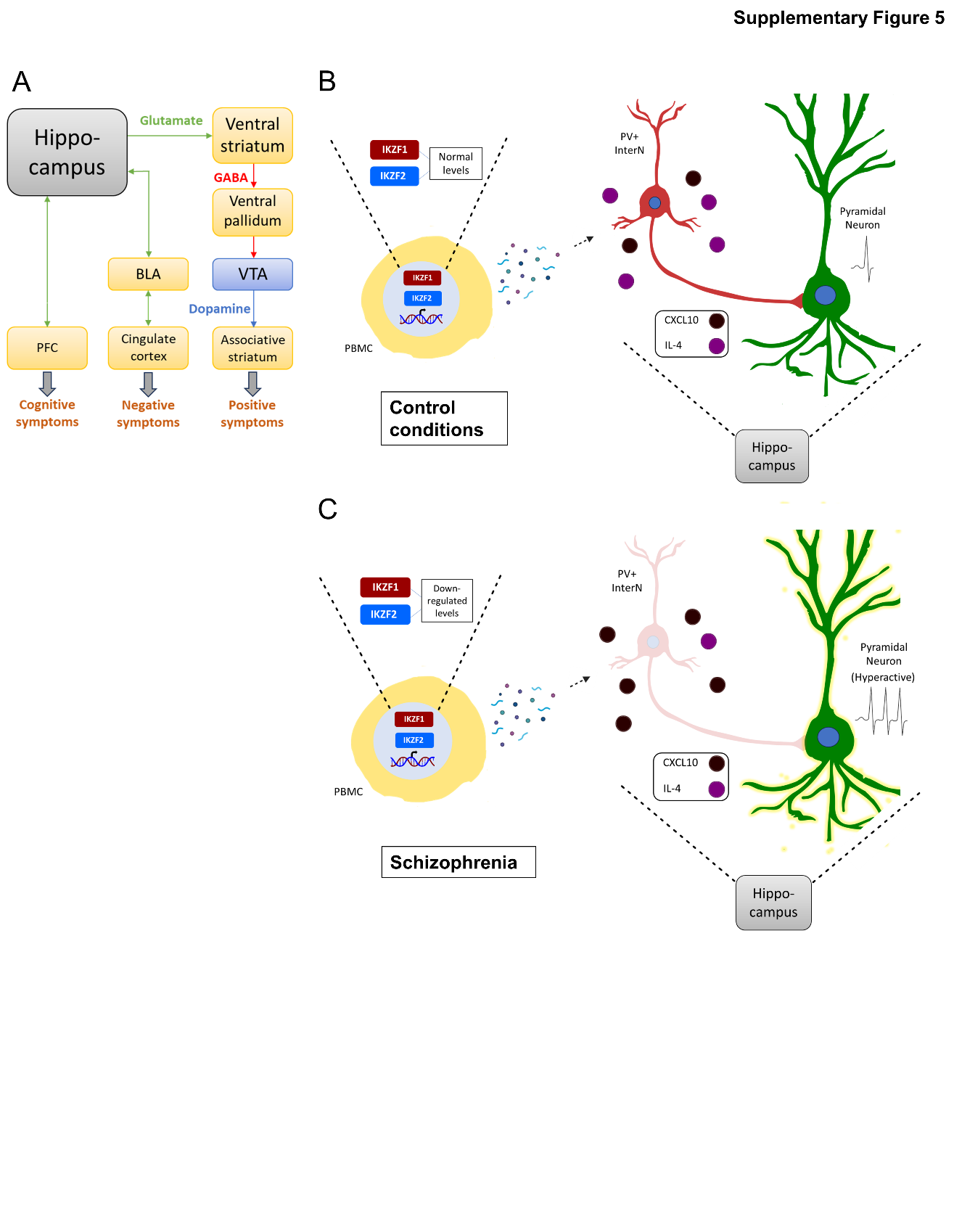

**Supplementary figure 7. Proposed model**. (**a**) In the context of schizophrenia, the hyperactive and dysrhythmic hippocampus is proposed to be associated with the appearance of the three categories of symptoms (positive, negative, and cognitive). The overdrive in the responsivity of dopaminergic neurons in the ventral tegmental area (VTA) that project to the associative striatum is proposed to underlie the positive symptoms. Besides, the hyperactive hippocampus can also be implicated in the dysfunction of other circuits. The hippocampus-basolateral amygdala (BLA) connection would interfere with the BLA-limbic cortical pathway implicated in the control of emotional responses, probably contributing to the appearance of negative symptoms. The hippocampus-PFC projection would possibly alter the PFC activity and rhythmicity, generating dysfunctions at the cognitive level. Taking in account our results, (**b**-**c**) the proposed model to explain the reduced quantity of parvalbumin-positive interneurons (PV+) in the context of schizophrenia would be through the altered secretome of the PBMCs with downregulated levels of both, *IKZF1* and *IKZF2*. The dysregulated levels of CXCL10 and Il-4 would be underlying the diminished number (or function) of PV+ interneurons. The reduced inhibition generated by the lack of these GABAergic PV+ interneurons would lead to the hyperactive and dysrhythmic state in the hippocampus and this in turn contributing to the appearance of all types of symptoms of schizophrenia.

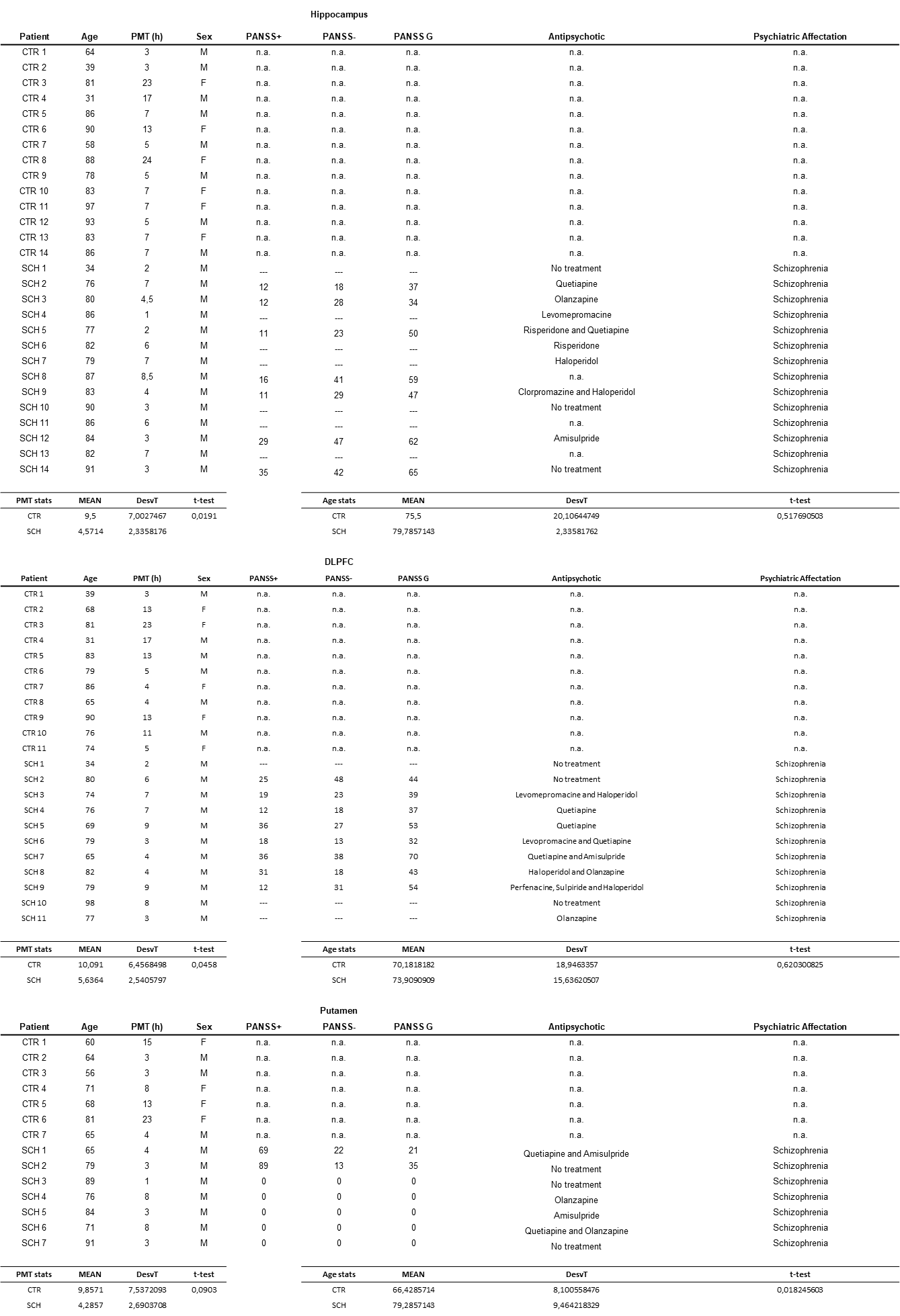

**Supplementary table 1**. Demographic characteristics from the patients of whom we obtained the post-mortem tissue including the hippocampus, the dorsolateral prefrontal cortex (DLPFC) and Putamen. MPT: Post-mortem time. Scores from the positive (PANSS+), negative (PANSS-) and general (PANSS G) are depicted from PANSS scale. CTR: Control patients, SCH: Patients with schizophrenia. Data are mean ± DesvT and they were analyzed using the two-tailed Student t-test.

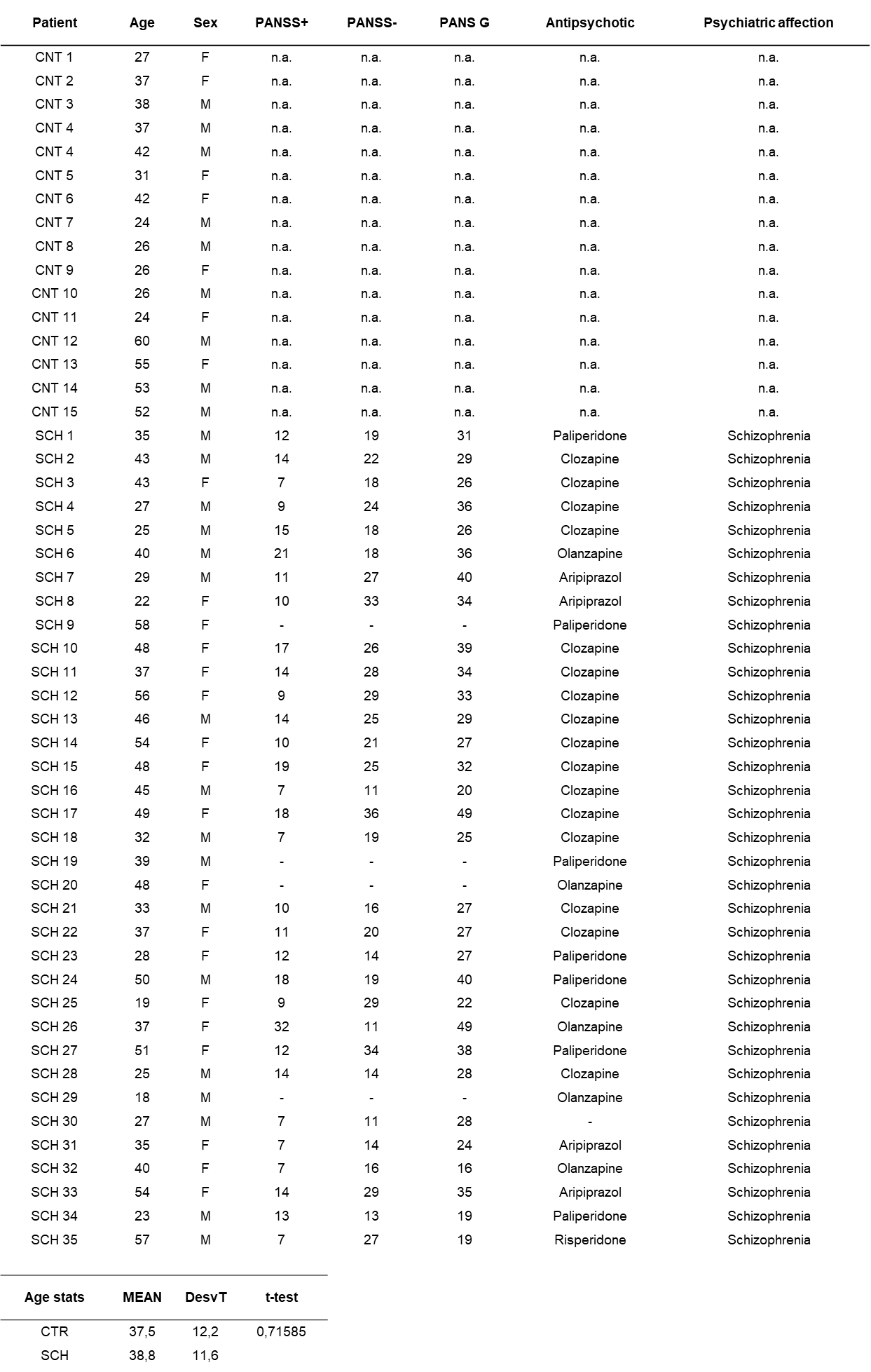

**Supplementary table 2**. Demographic characteristics from the patients of whom we obtained the peripheral blood mononuclear cells (PBMCs) supernatants. CTR: Control patients; SCH: Patients with schizophrenia. Data are mean ± DesvT and they were analyzed using the two-tailed Student t-test.

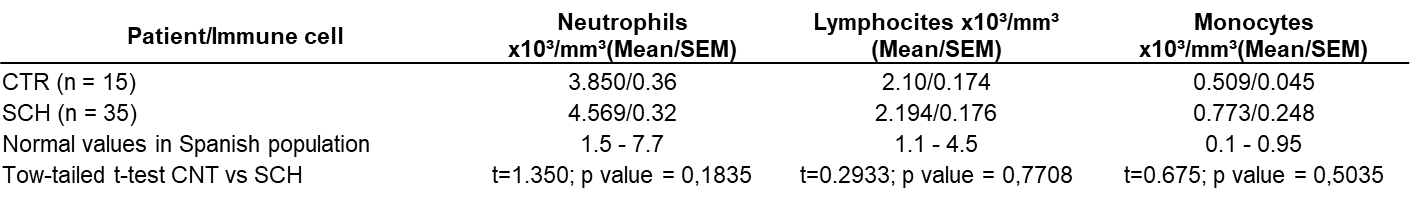

**Supplementary table 3**. Descriptive statistics of immune circulating cells in blood samples from controls (CTR) and patients with schizophrenia (SCH) was conducted in the current study. Mean values of circulating neutrophils, lymphocytes, and monocytes densities from blood samples are depicted. No differences were observed between the groups in our cohort. Densities were found to be within normal distributions compared to the Spanish population. Data are mean ± SEM and they were analyzed using the two-tailed Student t-test.

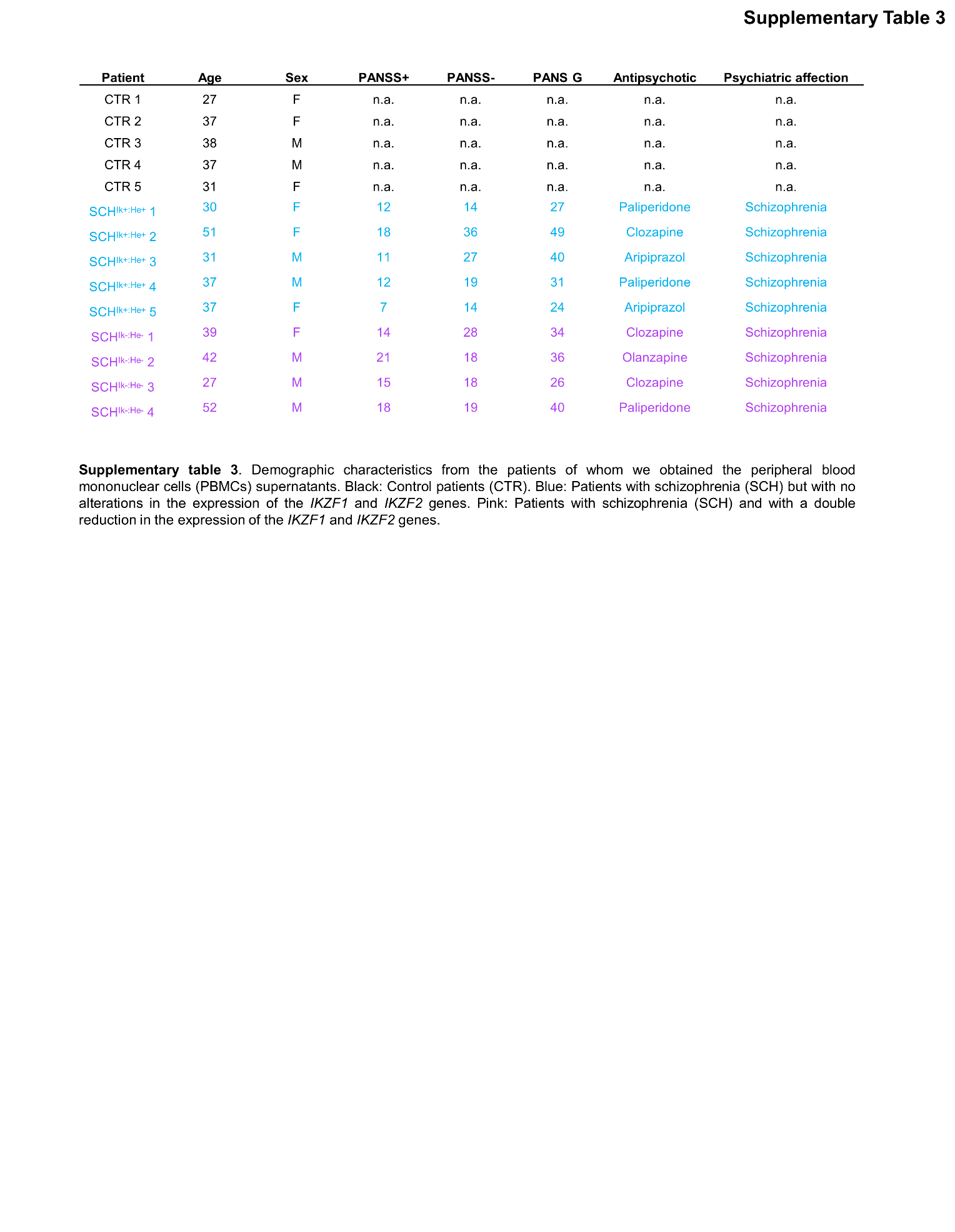

**Supplementary table 4**. Demographic characteristics from the patients of whom we obtained the peripheral blood mononuclear cells (PBMCs) supernatants. Black: Control patients (CTR). Blue: Patients with schizophrenia (SCH) but with no alterations in the expression of the *IKZF1* and *IKZF2* genes. Pink: Patients with schizophrenia (SCH) and with a double reduction in the expression of the *IKZF1* and *IKZF2* genes.
